# Combinatorial Control through Allostery

**DOI:** 10.1101/508226

**Authors:** Vahe Galstyan, Luke Funk, Tal Einav, Rob Phillips

## Abstract

Many instances of cellular signaling and transcriptional regulation involve switch-like molecular responses to the presence or absence of input ligands. To understand how these responses come about and how they can be harnessed, we develop a statistical mechanical model to characterize the types of Boolean logic that can arise from allosteric molecules following the Monod-Wyman-Changeux (MWC) model. Building upon previous work, we show how an allosteric molecule regulated by two inputs can elicit AND, OR, NAND and NOR responses, but is unable to realize XOR or XNOR gates. Next, we demonstrate the ability of an MWC molecule to perform ratiometric sensing - a response behavior where activity depends monotonically on the ratio of ligand concentrations. We then extend our analysis to more general schemes of combinatorial control involving either additional binding sites for the two ligands or an additional third ligand and show how these additions can cause a switch in the logic behavior of the molecule. Overall, our results demonstrate the wide variety of control schemes that biological systems can implement using simple mechanisms.

## Introduction

A hallmark of cellular signaling and regulation is combinatorial control. Disparate examples ranging from metabolic enzymes to actin polymerization to transcriptional regulation involve multiple inputs that often give rise to a much richer response than what could be achieved through a single-input. For example, the bacterial enzyme phosphofructokinase in the glycolysis pathway is allosterically regulated by both ADP and PEP.^1^ Whereas PEP serves as an allosteric inhibitor, ADP is both an allosteric activator and a competitive inhibitor depending upon its concentration. This modulation by multiple allosteric ligands gives rise to a complex control of the flux through the glycolytic pathway: increasing ADP concentration first increases the activity of phosphofructokinase (via the allosteric modulation) but ultimately decreases it (from competitive inhibition). Another example is offered by the polymerization of actin at the leading edge of motile cells. In particular, the presence of two ligands, Cdc42 and PIP2, is required to activate the protein N-WASP by binding to it in a way that permits it to then activate the Arp2/3 complex and stimulate actin polymerization.^2^

In the context of transcriptional regulation, an elegant earlier work explored the conditions under which transcriptional regulatory networks could give rise to the familiar Boolean logic operations, like those shown in Figure 1.^3^ There it was found that the combined effect of two distinct transcription factors on the transcriptional activity of a given promoter depend upon their respective binding strengths as well as the cooperative interactions between each other and the RNA polymerase. Indeed, by tuning the binding strengths and cooperativity parameters, one could generate a panoply of different logic gates such as the familiar AND, OR, NAND (NOT-AND) and NOR (NOT-OR) gates, known from the world of digital electronics.^3^

**Figure 1.**
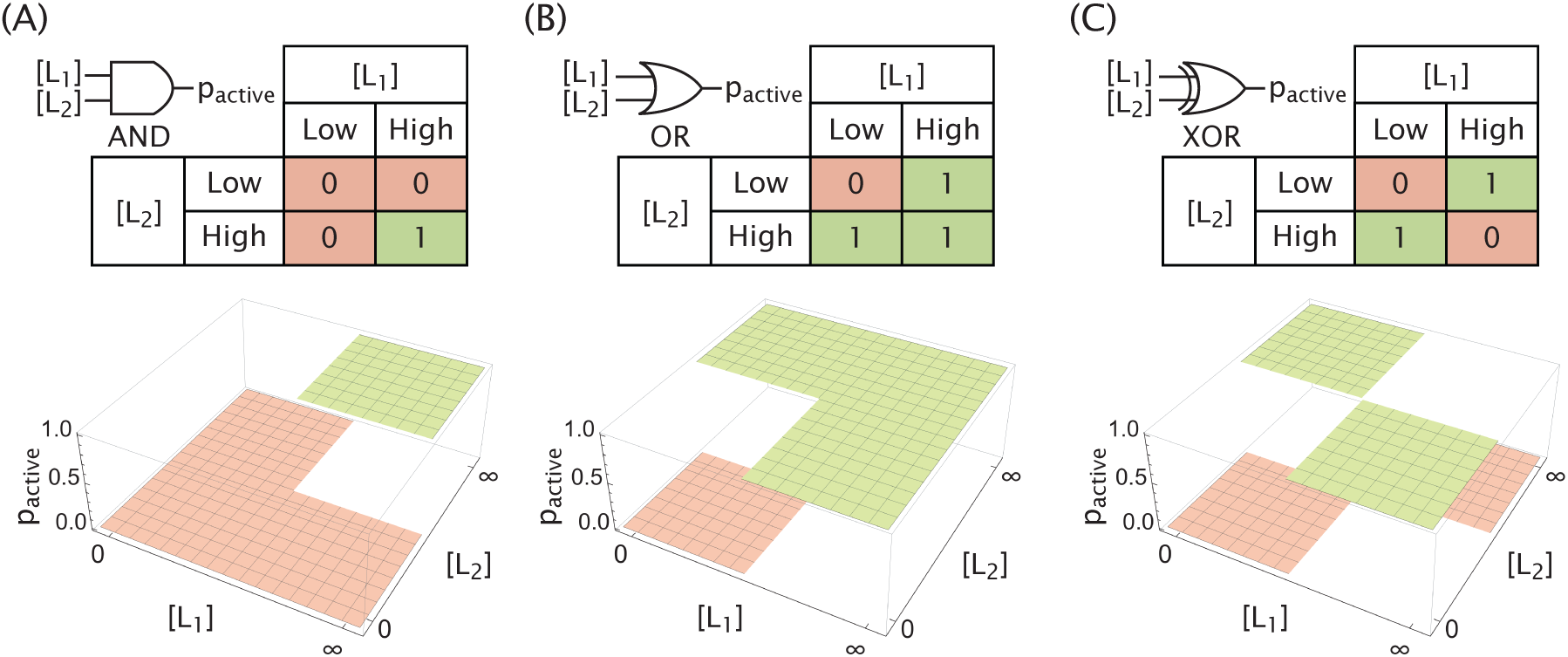
Logic gates as molecular responses. The (A) AND, (B) OR, and (C) XOR gates are represented through their corresponding logic tables as well as target activity profiles regulated by two ligands. The behavior of each gate is measured solely by its activity in the absence and at saturating concentrations of each ligand and not by the character of the active/inactive transition.

Here we explore the diversity of combinatorial responses that can be effected by a single allosteric molecule by asking if such molecules can yield multi-input combinatorial control in the same way that transcriptional networks have already been shown to. Specifically, we build on earlier work that shows that an allosteric molecule described by the Monod-Wyman-Changeux (MWC) model can deliver input-output functions similar to the ideal logic gates described in Figure 1.^4–6^ In the MWC model, an allosteric molecule exists in a thermodynamic equilibrium between active and inactive states, with the relative occupancy of each state being modulated by regulatory ligands.^7^ We use statistical mechanics to characterize the input-output response of such a molecule in the limits where each of the two ligands is either absent or at a saturating concentration and determine the necessary conditions to form the various logic gates, with our original contribution on this point focusing on a systematic exploration of the MWC parameter space for each logic gate.

We then analyze the MWC response modulated by two input ligands but outside of traditional Boolean logic functions. In particular, we show how, by tuning the MWC parameters, the response (probability of the allosteric protein being active) in any three of the four concentration limits can be explicitly controlled, along with the ligand concentrations at which transitions between these limit responses occur. Focusing next on the profile of the response near the transition concentrations, we demonstrate how an MWC molecule can exhibit ratiometric sensing which was observed experimentally in the bone morphogenetic protein (BMP) signaling pathway^8^ as well as in galactose metabolic (GAL) gene induction in yeast.^9^

Additionally, we extend our analysis of logic responses to cases beyond two-ligand control with a single binding site for each ligand. We first discuss the effect of the number of binding sites on the logic response and demonstrate how altering that number, which can occur through evolution or synthetic design, is able to cause a switch in the logic-behavior of an MWC molecule, such as transitioning from AND into OR behavior. Next, we explore the increased diversity of logic responses that can be achieved by three-ligand MWC molecules compared with the two-ligand case and offer an interesting perspective on the role of the third ligand as a regulator that can switch the logic-behavior formed by the other two ligands. We end by a discussion of our theoretical results in the context of a growing body of experimental works on natural and *de novo* designed molecular logic gates. In total, these results hint at simple mechanisms that biological systems can utilize to refine their combinatorial control.

## Results

### Logic Response of an Allosteric Protein Modulated by Two Ligands

Consider an MWC molecule, as shown in Figure 2, that fluctuates between active and inactive states (with ∆*ε*_AI_ defined as the free energy difference between the inactive and active states in the absence of ligand). We enumerate the entire set of allowed states of activity and ligand occupancy, along with their corresponding statistical weights. The probability that this protein is active depends on the concentrations of two input molecules, [L_1_] and [L_2_], and is given by

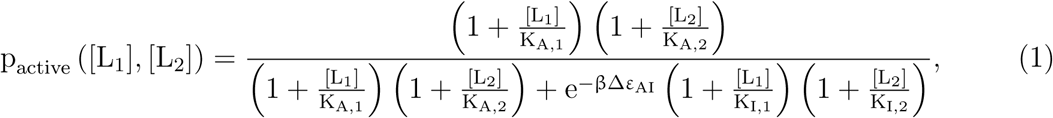

**Figure 2.**
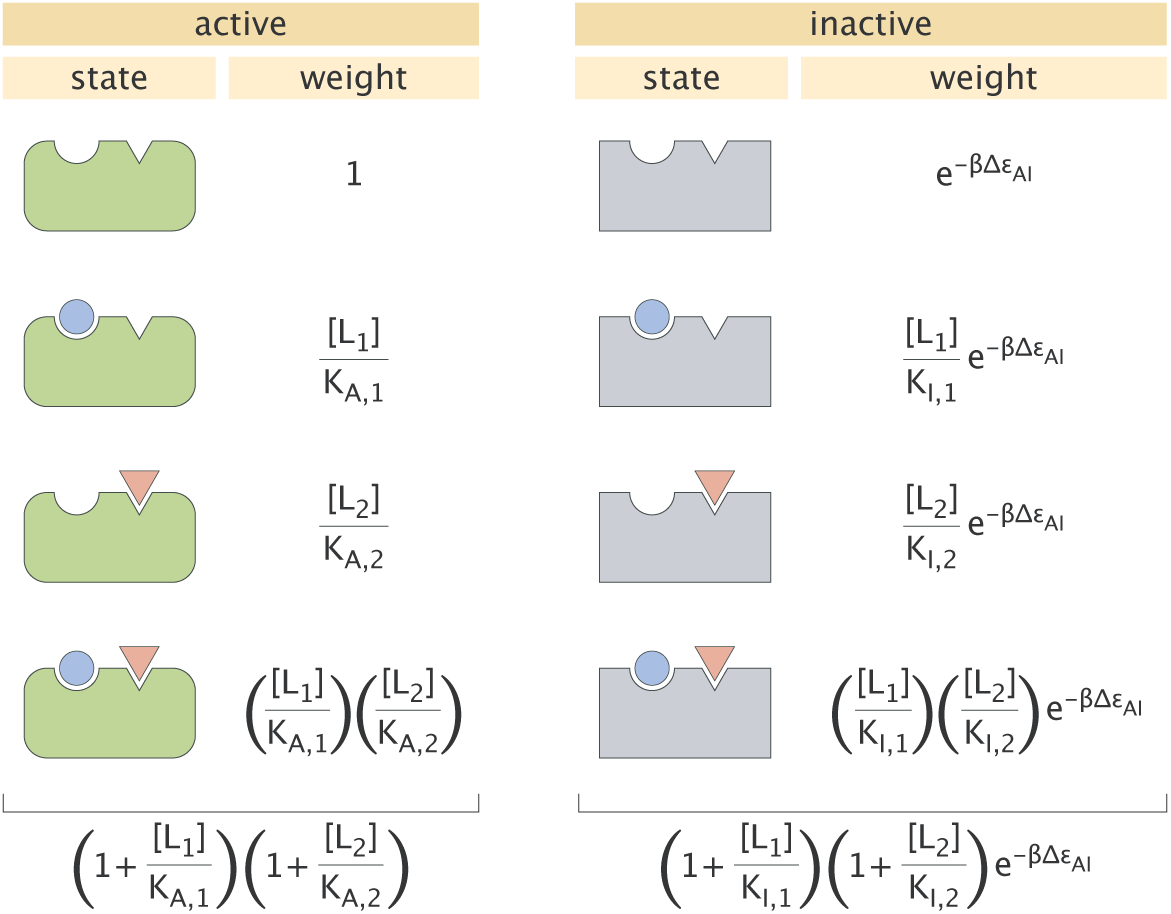
States and weights for the allosteric protein. The two different ligands (blue circle (i = 1) and red triangle (i = 2)) are present at concentrations [L_i_] and with a dissociation constant K_A,i_ in the active state and K_I,i_ in the inactive state. The energetic difference between the inactive and active states is denoted by ∆*ε*_AI_ = *ε*_I_ – *ε*_A_. Total weights of the active and inactive states are shown below each column and are obtained by summing all the weights in that column.

where K_A,i_ and K_I,i_ are the dissociation constants between the i^th^ ligand and the active or inactive protein, respectively. We begin with the two-input case such that i = 1 or 2.

To determine whether this allosteric protein can serve as a molecular logic gate, we first evaluate the probability that it is active when each ligand is either absent ([L_i_] → 0) or at a saturating concentration ([L_i_] → *∞*). Figure 3A evaluates these limits for eq 1, where we have introduced the parameters 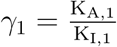 and 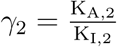 to simplify the results.

**Figure 3.**
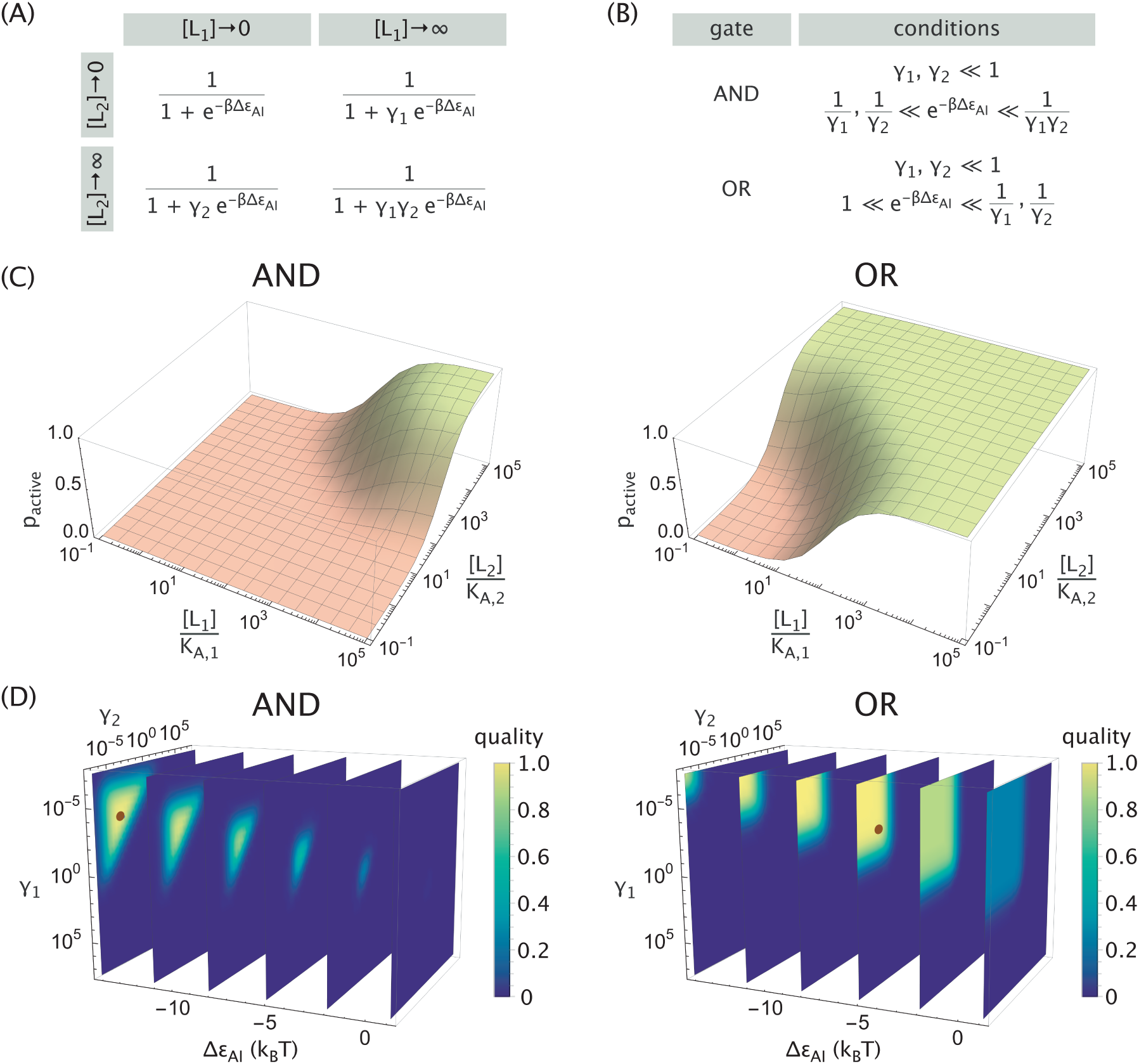
Logic gate realization of an allosteric protein with two ligands. (A) Probability that the protein is active (p_active_) in different limits (rows and columns of the matrix) of ligand concentrations, where 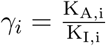. (B) Conditions on the parameters that lead to an AND or OR response. (C) Realizations of the AND and OR logic gates. Parameters used were K_A,1_ = K_A,2_ = 2.5 × 10^−8^ M, K_I,1_ = K_I,2_ = 1.5 × 10^−4^ M, and ∆*ε*_AI_ = −14.2 k_B_T for the AND gate or ∆*ε*_AI_ = 5.0 k_B_T for the OR gate. (D) Quality of AND (eq 3) and OR (eq 4) gates across parameter space. The brown dots indicate the high quality gates in Panel C.

The probabilities in Figure 3A can be compared to the target functions in Figure 1 to determine the conditions on each parameter that would be required to form a given logic gate. For example, the AND, OR, and XOR gates require that in the absence of either ligand ([L_1_] = [L_2_] = 0), there should be as little activity as possible, thereby requiring that the active state has a higher (more unfavored) free energy than the inactive state (e^−β∆*ε*^^AI^ ≫ 1). We note that in the context of transcriptional regulation, this limit of activity in the absence of ligands is called the leakiness, ^10^ and it is one of the distinguishing features of the MWC model in comparison with other allosteric models such as the Koshland-Némethy-Filmer (KNF) model that exhibits no leakiness.

For the AND and OR gates, the condition that p_active_ ≈ 1 when both ligands are saturating ([L_1_], [L_2_] → ∞) requires that *γ*_1_*γ*_2_e^−β∆*ε*^^AI^ ≪ 1. The two limits where one ligand is absent while the other ligand is saturating lead to the conditions shown in Figure 3B for the AND and OR gates, with representative response profiles shown in Figure 3C using parameter values from the single-ligand allosteric nicotinic acetylcholine receptor.^11^ We relegate the derivations to Appendix A, where we also demonstrate that the XOR gate cannot be realized with the form of p_active_ in eq 1 unless explicit cooperativity is added to the MWC model. In addition, we show that the NAND, NOR, and XNOR gates can be formed if and only if their complementary AND, OR, and XOR gates can be formed, respectively, by replacing ∆*ε*_AI_ → −∆*ε*_AI_ and 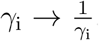. Finally, Figure 3C demonstrates that the same dissociation constants K_A,i_ and K_I,i_ can give rise to either AND or OR behavior by modulating ∆*ε*_AI_, with the transition between these two logic gates occurring at 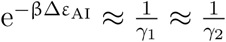 (this corresponds to ∆*ε*_AI_ ≈ −9 k_B_T for the values of K_A,i_ and K_I,i_ in Figure 3).

To explore the gating behavior changes across parameter space, we define a quality metric for how closely p_active_ matches its target value at different concentration limits for a given idealized logic gate,

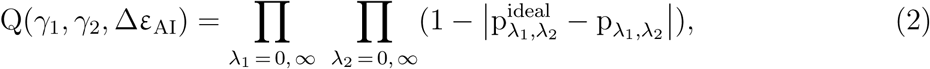

where p_λ1,λ2_ = p_active_ ([L_1_] → λ_1_, [L_2_] → λ_2_). A value of 1 (high quality gate) implies a perfect match between the target function and the behavior of the allosteric molecule while a value near 0 (low quality gate) suggests that the response behavior deviates from the target function in at least one limit.

From eq 2, the quality for the AND gate becomes

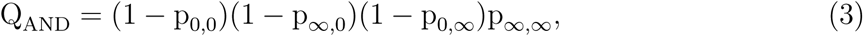

while for the OR gate it takes on the form

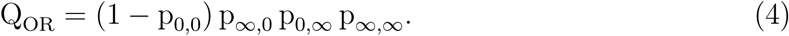

Figure 3D shows the regions in parameter space where the protein exhibits these gating behaviors (the high quality gates from Figure 3C are denoted by brown dots). More specifically, for a fixed ∆*ε*_AI_, the AND behavior is achieved in a finite triangular region in the *γ*_1_-*γ*_2_ plane which grows larger as ∆*ε*_AI_ decreases. The OR gate, on the other hand, is achieved in an infinite region defined by *γ*_1_, *γ*_2_ ≲ e^β∆*ε*^^AI^. In either case, a high quality gate can be obtained only when the base activity is very low (∆*ε*_AI_ ≲ 0) and when both ligands are strong activators (*γ*_1_, *γ*_2_ ≪ 1), in agreement with the derived conditions (Figure 3B). Lastly, we note that the quality metrics for AND/OR and their complementary NAND/NOR gates obey a simple relation, namely, Q_AND/OR_ (*γ*_1_, *γ*_2_*, ∆*ε**_AI_) = Q_NAND/NOR_ (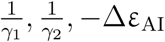), which follows from the functional form of eq 2 and the symmetry between the two gates (see Appendix A).

## General Two-Ligand MWC Response

We next relax the constraint that p_active_ must either approach 0 or 1 in the limits of no ligand or saturating ligand and consider the general behavior that can be achieved by an MWC molecule in the four limits shown in Figure 3A. Manipulating the three parameters (*γ*_1_, *γ*_2_ and ∆*ε*_AI_) enables us to fix three of the four limits of p_active_, and these three choices determine the remaining limit. For example, the parameters in Figure 4A were chosen so that p_0,0_ = 0.5 (∆*ε*_AI_ = 0), p_0,∞_ ≈ 0.9 (*γ*_2_ = 0.1), and p_∞,0_ ≈ 0.05 (*γ*_1_ = 20), which fixed p_∞,∞_ ≈ 0.3 for the final limit.

**Figure 4.**
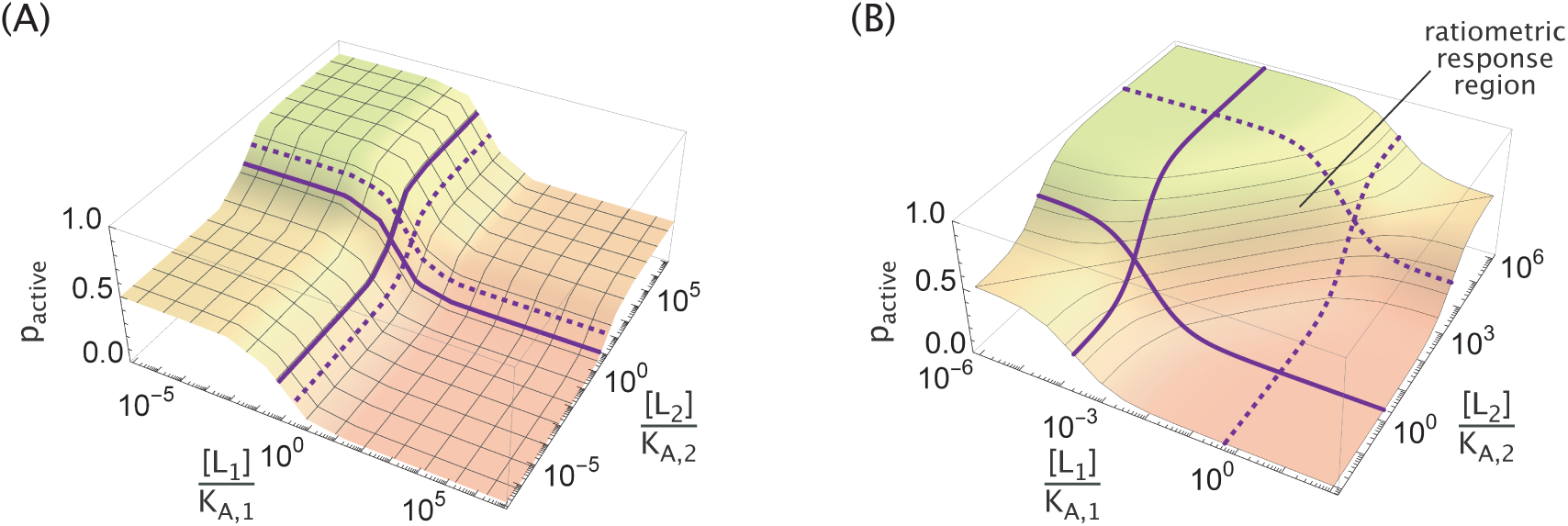
General MWC response with two ligands. (A) Three of the four limits of ligand concentrations ([L_1_], [L_2_] → 0 or ∞) can be fixed by the parameters ∆*ε*_AI_, *γ*_1_, and *γ*_2_. Additionally, the midpoint of the [L_i_] response when [L_j_] → 0 (solid purple curve) or [L_j_] →∞ (dashed purple curve) can be adjusted. (B) Within the region determined by the four midpoints, the MWC response becomes ratiometric ^8^ where the concentration ratio of the two ligands determines the activity of the molecule. This is illustrated by the diagonal contour lines of constant p_active_ in the ratiometric response region.

In addition to the limits of p_active_, the locations of the transitions between these limits can be controlled by changing K_A,i_ and K_I,i_ while keeping 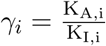 constant. In Appendix B we generalize previous results for the transition of a single-ligand MWC receptor^12^ to the present case of two ligands. Interestingly, we find that the midpoint 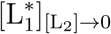 of the response in the absence of [L_2_] (solid curve in Figure 4A) is different from the midpoint 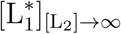 of the response at saturating [L_2_] (dashed curve in Figure 4A), with analogous statements holding for the second ligand. More precisely, the two transition points occur at

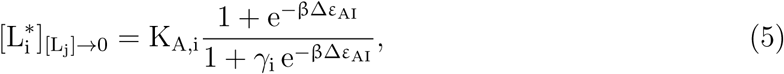

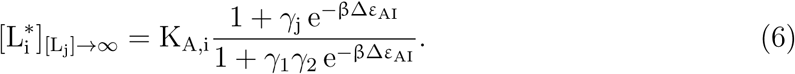

Notably, the ratio

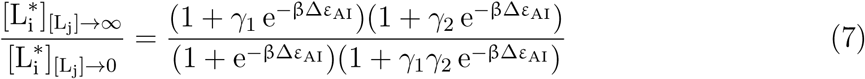

is invariant to ligand swapping (i ↔ j); hence, the transition zones, defined as the concentration intervals between solid and dotted curves, have identical sizes for the two ligands, as can be seen in Figure 4.

The MWC response has its steepest slope when the ligand concentration is within the range set by 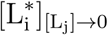 and 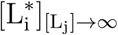, and interesting response behaviors can arise when both ligand concentrations fall into this regime. For example, Antebi *et al.* recently showed that the BMP pathway exhibits ratiometric response where pathway activity depends monotonically on the ratio of the ligand concentrations. ^8^ Similar response functions have also been observed in the GAL pathway in yeast, where gene induction is sensitive to the ratio of galactose and glucose.^9^ Such behavior can be achieved within the highly sensitive region of the MWC model using one repressor ligand (L_1_) and one activator ligand (L_2_), as shown in Figure 4B. Parameters chosen for demonstration are ∆*ε*_AI_ = 0, K_A,1_ = K_A,2_ and 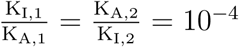. In this regime, the probability of the protein being active gets reduced to

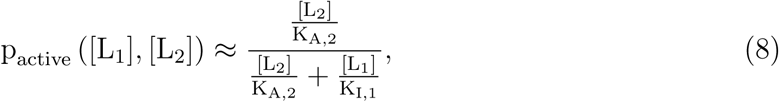

which clearly depends monotonically on the [L_2_]*/*[L_1_] ratio (see Appendix B for details). We note that the region over which the ratiometric behavior is observed can be made arbitrarily large by decreasing the ratios 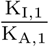and 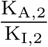

### Modulation by Multiple Ligands

A much richer repertoire of signaling responses is available to an MWC protein if we go beyond two ligand inputs with a single binding site for each, as exhibited by phosphofructokinase, for example. Though earlier we mentioned phosphofructokinase in the context of two of its input ligands, in fact, this enzyme has even more inputs than that and thus provides a rich example of multi-ligand combinatorial control. ^1^ To start exploring the diversity of these responses, we generalize eq 1 to consider cases with N input ligands, where the i^th^ ligand has n_i_ binding sites, concentration [L_i_], and dissociation constants K_A,i_ and K_I,i_ with the molecule’s active and inactive states, respectively. In general, it is impractical to write the states and weights as we have done in Figure 2, since the total number of possible states, given by 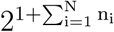, grows exponentially with the number of binding sites. However, by analogy with the earlier simple case, the general formula for the probability that the protein is active can be written as

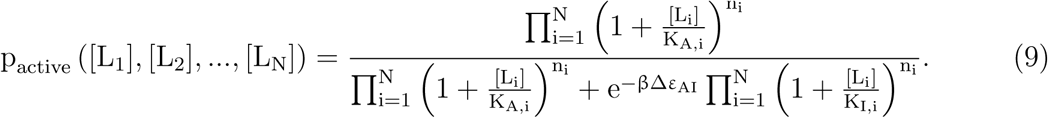

We first consider an MWC molecule with N = 2 input ligands as in the previous section but with n_i_ ligand binding sites for ligand i. As derived in Appendix C, the criteria for the AND and OR gates are identical to those for a protein with n_i_ = 1 binding site per ligand, except that we make the 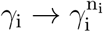 substitution in the conditions shown in Figure 3B. The protein thus exhibits OR behavior if 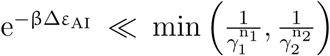 or AND behavior if 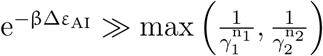.

Over evolutionary time or through synthetic approaches, the number of binding sites displayed by a single molecule can be tuned, enabling such systems to test a variety of responses with a limited repertoire of regulatory molecules. Since *γ*_1_, *γ*_2_ ≪ 1, increasing the number of binding sites while keeping all other parameters the same can shift a response from AND→OR as shown in Figure 5. The opposite logic switching (OR→AND) is similarly possible by decreasing the number of binding sites, and analogous results can be derived for the complementary NAND and NOR gates (see Appendix C). In the limit where the number of binding sites becomes large (n_1_, n_2_ ≫ 1), an allosteric molecule’s behavior will necessarily collapse into OR logic provided *γ*_1_, *γ*_2_ < 1, since the presence of either ligand occupying the numerous binding sites has sufficient free energy to overcome the active-inactive free energy difference ∆*ε*_AI_. In addition, having a large number of binding sites makes the p_active_ response sharper (Figure 5B), as has been seen in the context of chromatin remodeling where ~150 bp of DNA “buried” within a nucleosome can be made available for transcription by the binding of multiple transcription factors. ^13^

**Figure 5.**
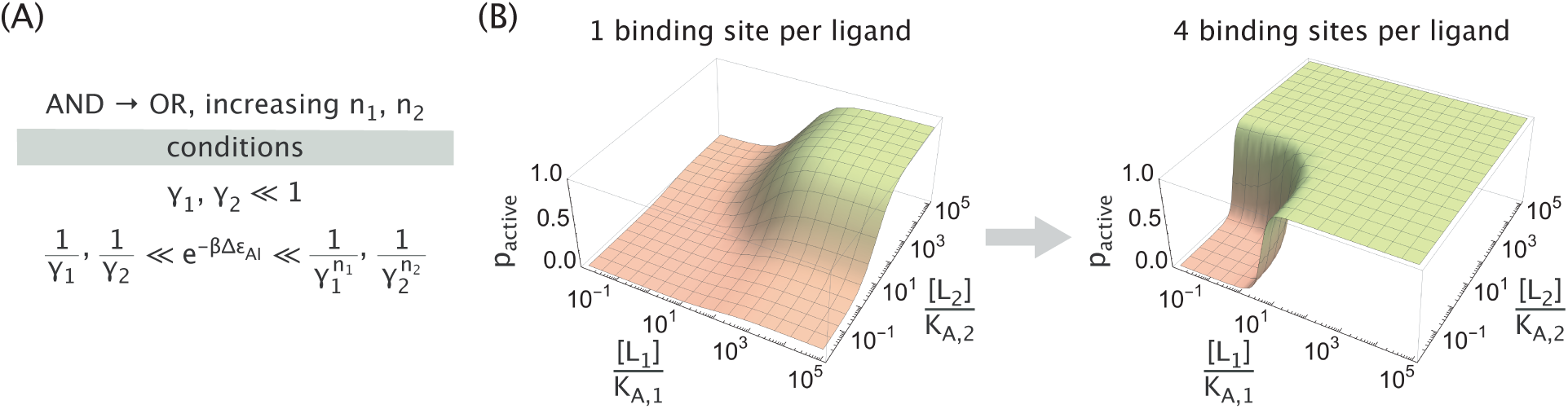
Increased number of binding sites can switch the logic of an MWC protein from AND into OR. (A) Parameter conditions required for AND → OR switching upon an increase in the number of binding sites. (B) Representative activity plots showing the AND → OR switching. Parameters used were K_A,i_ = 2.5 × 10^−8^ M, K_I,i_ = 2.5 × 10^−6^ M and ∆*ε*_AI_ = −7 k_B_T.

Next, we examine an alternative possibility of generalizing the MWC response, namely, considering a molecule with N = 3 distinct ligands, each having a single binding site (n_i_ = 1). The logic response is now described by a 2 × 2 × 2 cube corresponding to the activity at low and saturating concentrations of each of the three ligands (an example realization is shown in Figure 6A). Since each of the 8 cube elements can be either OFF or ON (red and green circles, respectively), the total number of possible responses becomes 2^8^ = 256. This number, however, includes functionally redundant responses, as well as ones that are not admissible in the MWC framework. We therefore eliminate these cases in order to accurately quantify the functional diversity of 3-input MWC proteins.

**Figure 6.**
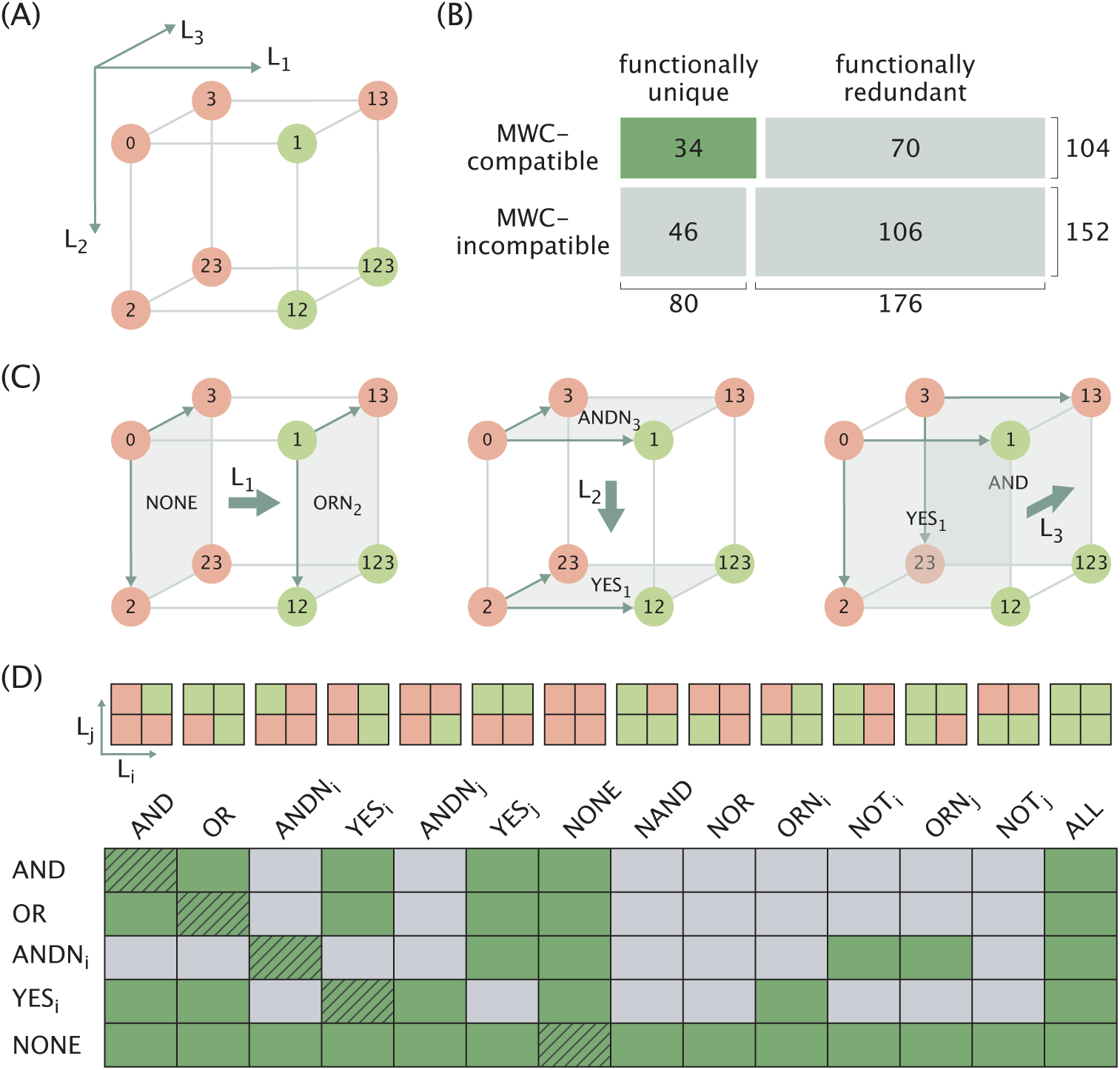
Third ligand expands the combinatorial diversity of logic responses and enables logic switching. (A) Cubic diagram of a representative molecular logic response. The label “0” stands for the limit when all ligands are at low concentrations. Each digit in the labels of other limits indicates the high concentration of the corresponding ligand (for example, in the “12” limit the ligands 1 and 2 are at high concentrations). Red and green colors indicate the OFF and ON states of the molecule, respectively. (B) Diagram representing the numbers of 3-ligand logic gates categorized by their MWC compatibility and functional uniqueness. The area of each cell is proportional to the number of gates in the corresponding category. (C) Demonstration of different logic transitions induced by a third ligand (thick arrows) on the example of the 3-input gate in Panel A. (D) Table of all possible logic transitions (row → column, green cells) inducible by a third ligand in the MWC framework. Schematics of the 14 MWC-compatible 2-ligand gates corresponding to each column entry are displayed on top (i and j represent different ligands). Results for the transitions between logical complements (NOT row → NOT column) are identical to the results for row → column transitions and are not shown. Trivial transitions between identical gates where the third ligand has no effect are marked with hatching lines.

We consider two responses to be functionally identical if one can be obtained from another by relabeling the ligands, e.g. (1, 2, 3) → (3, 1, 2). Eliminating all redundant responses leaves 80 unique cases out of the 256 possibilities (see Appendix D). In addition, since the molecule’s activity in the eight ligand concentration limits is determined by only four MWC parameters, namely, {∆*ε*_AI_, *γ*_1_, *γ*_2_, *γ*_3_}, we expect the space of possible 3-input gates to be constrained (analogous to XOR/XNOR gates being inaccessible to 2-input MWC proteins). Imposing the constraints leaves 34 functionally unique logic responses that are compatible with the MWC framework (see Figure 6B for the summary statistics and Appendix D for the detailed discussion of how the constraints were imposed).

In addition to expanding the scope of combinatorial control relative to the two-input case, we can think of the role of the third ligand as a regulator whose presence switches the logic performed by the other two ligands. We illustrate this role in Figure 6C by first focusing on the leftmost cubic diagram. The gating behavior on the left face of the cube (in the absence of L_1_) exhibits NONE logic while the behavior on the right face of the cube (in the presence of saturating L_1_) is the ORN_2_ logic (see the schematics at the top of Figure 6D for the definition of all possible gates). In this way, adding L_1_ switches the logic of the remaining two ligands from NONE → ORN_2_. In a similar vein, adding L_2_ changes the logic from ANDN_3_ → YES_1_, while adding L_3_ causes a YES_1_ → AND switch.

We repeat the same procedure for all functionally unique 3-ligand MWC gates (see Appendix D) and obtain a table of all possible logic switches that can be induced by a third ligand (green cells in Figure 6D that indicate row → column logic switches). As we can see, a large set of logic switches are feasible, the majority of which (the left half of the table) do not involve a change in the base activity (i.e., activity in the absence of the two ligands). Comparatively fewer transitions that involve flipping of the base activity from OFF to ON are possible (the right half of the table).

As a demonstration of the regulatory function of the third ligand, we show two examples of logic switching induced by increasing [L_3_], namely, AND→OR (Figure 7A,B) and AND→YES_1_ (Figure 7C,D), along with the parameter conditions that need to be satisfied to enable such transitions (see Appendix D for derivations). An interesting perspective is to view the L_3_ ligand as a modulator of the free energy difference ∆*ε*_AI_. For example, when [L_3_] = 0, the protein behaves identically to the N = 2 case given by eq 1; at a saturating concentration of L_3_, however, the protein behaves as if it had N = 2 ligands with a modified free energy difference 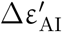 given by

**Figure 7.**
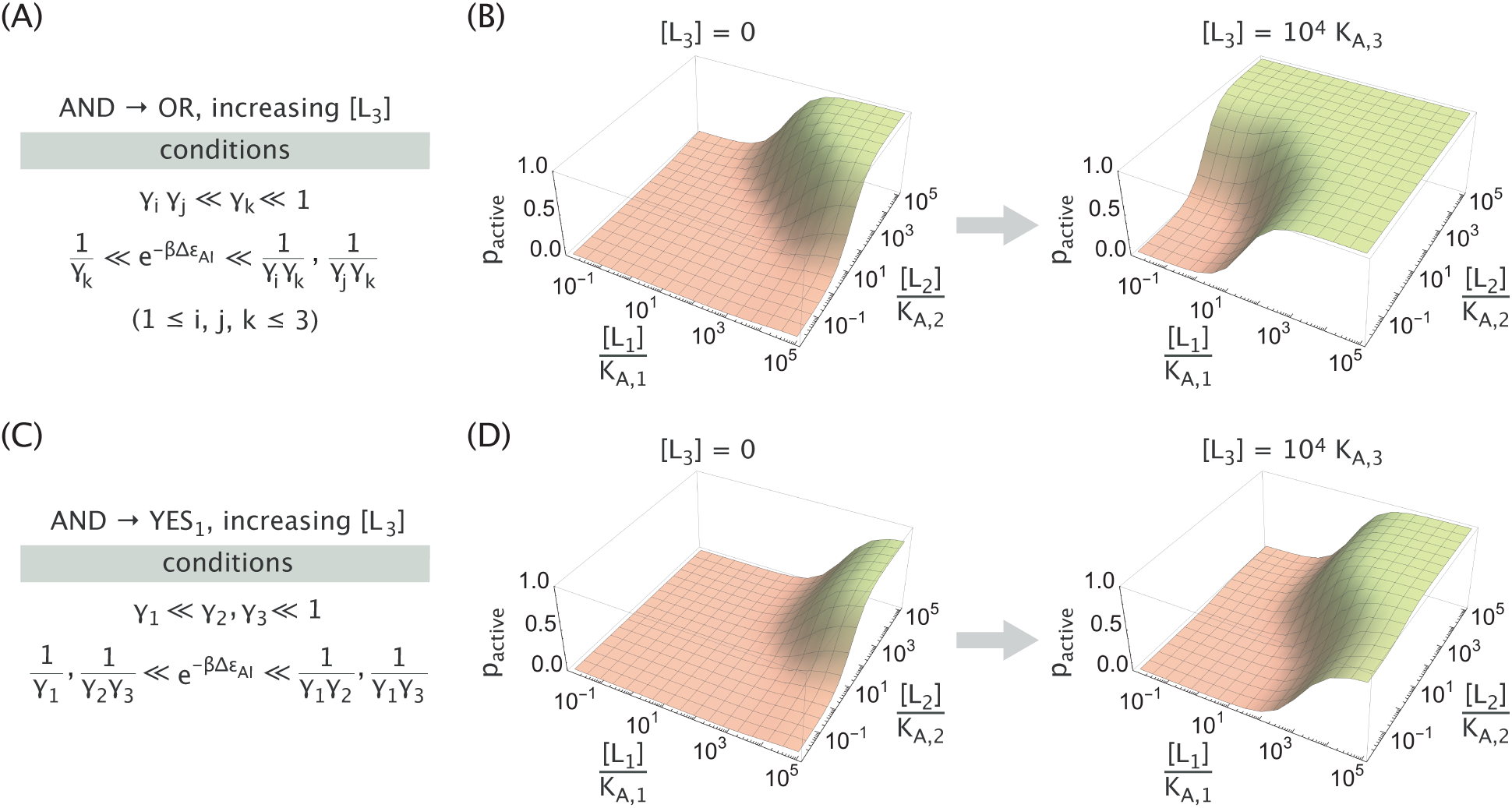
Example logic switches induced by the third ligand. Parameter conditions and representative activity plots of an allosteric molecule exhibiting AND logic in the absence of the third ligand, while exhibiting OR logic (A,B) or YES_1_ logic (C,D) when L_3_ is present at a saturating concentration. Parameters used were K_A,i_ = 2.5 × 10^−8^ M and K_I,i_ = 2.5 × 10^−4^ M in Panel B, K_A,i_ = 2.5 × 10^−8^ M, K_I,1_ = 2.5 × 10^−4^ M and K_I,2/3_ = 2.5 × 10^−6^ M in panel D, along with ∆*ε*_AI_ = −12 k_B_T in both panels.

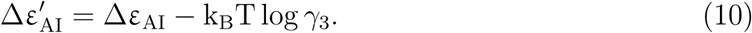

From this perspective, the third ligand increases the effective free energy difference in the examples shown in Figure 7, since in both cases the *γ*_3_ ≪ 1 condition is satisfied. For the AND→OR transition, the increase in ∆*ε*_AI_ is sufficient to let either of the two ligands activate the molecule (hence, the OR gate). In the AND→YES_1_ transition, the change in ∆*ε*_AI_ utilizes the asymmetry between the binding strengths of the two ligands (*γ*_1_ ≫ *γ*_2_) to effectively “silence” the activity of the ligand L_2_. We note in passing that such behavior for the N = 3 allosteric molecule is reminiscent of a transistor which can switch an input signal in electronics.

## Discussion and Conclusions

Combinatorial control is a ubiquitous strategy employed by cells. Networks of cellular systems of different kinds, such as transcriptional, ^14,15^ signaling,^16^ or metabolic,^1^ integrate information from multiple inputs in order to produce a single output. The statistical mechanical MWC model we employ allows us to systematically explore the combinatorial diversity of output responses available to such networks and determine the conditions that the MWC parameters need to satisfy to realize a particular response.

In this paper, we built on earlier work to show that the response of an allosteric MWC molecule can mimic Boolean logic. Specifically, we demonstrated that a protein that binds to two ligands can exhibit an AND, OR, NAND, or NOR response (also shown by others^4–6^), where the former two cases require the protein to be inherently inactive and that both ligands preferentially bind to the active conformation, whereas the latter two cases require the converse conditions. We derived the MWC parameter ranges within which an allosteric protein would exhibit an AND or OR response (Figure 3B), and showed that the corresponding parameter ranges for NAND or NOR responses could be achieved by simply substituting 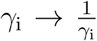 ∆*ε*_AI_ → – ∆*ε*_AI_ in the parameter condition equations (Appendix A.3). Since the NAND and NOR gates are known in digital electronics as universal logic gates, all other logic functions can be reproduced by hierarchically layering these gates. In the context of this work, such layering could be implemented if the MWC protein is an enzyme that only catalyzes in the active state so that its output (the amount of product) could serve as an input for the next enzyme, thereby producing more complex logic functions via allostery, though at the cost of noise amplification and response delays.

As in earlier work,^4,5^ we showed that the XOR and XNOR responses cannot be achieved within the original MWC framework (eq 1) but are possible when cooperativity between the two ligands is introduced (Appendix A.4). Biological XOR and XNOR behaviors are uncommon in non-transcriptional systems and have also been challenging for synthetic design and optimization.^17^ One of the few examples of such systems is a synthetic metallochromic chromophore whose transmittance output level is modulated by Ca^2+^ and H^+^ ions in a XOR-like manner.^18,19^

In addition to traditional Boolean logic, we recognized further manifestations of combinatorial control by two-ligand MWC proteins. In particular, we showed that the protein activity in three of the four ligand concentration limits can be set independently by tuning the MWC parameters *γ*_1_, *γ*_2_, and ∆*ε*_AI_, and that the ligand concentrations at which transitions between limit responses take place can be separately controlled by proportionally changing K_A,i_ and K_I,i_, while keeping 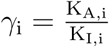 constant (eqs 5 and 6). We also showed that when the ranges of ligand concentrations are close to those transition values, then ratiometric sensing observed in the BMP^8^ and GAL pathways,^9^ can be recapitulated through the MWC model (Figure 4B), with larger regions of sensitivity achievable by an appropriate tuning of the parameters. We note that parameter “tuning” can be realized either through evolutionary processes over long time scales or synthetically, using mutagenesis or other approaches.^20^

Apart from altering the thermodynamic parameters such as the ligand binding affinity or the free energy of active and inactive protein conformations, the number of ligand binding sites of an allosteric molecule can also be changed. This can occur evolutionarily through recombination events, synthetically by engineering combinations of protein domains, ^21^ or through binding of competitive effectors that reduce the effective number of ligand binding sites. We found that these alterations in the number of ligand binding sites are capable of switching the logic behavior between AND↔OR or NAND↔NOR gates (Figure 5B). Since the MWC model has even been applied in unusual situations such as the packing of DNA into nucleosomes, ^13,22^ these results on combinatorial control can also be relevant for eukaryotic transcription. The opening of the nucleosome is itself often subject to combinatorial control because there can be multiple transcription factor binding sites within a given nucleosome, the number of which can also be tuned using synthetic approaches.^23–26^

Lastly, we generalized the analysis of logic responses for a molecule whose activity is modulated by three ligands, and identified 34 functionally unique and MWC-compatible gates out of 256 total possibilities. We offered a perspective on the function of any of the three ligands as a “regulator” that can cause a switch in the type of logic performed by the other two ligands and derived the full list of such switches (Figure 6D). Within the MWC model, the role of this regulatory ligand can be viewed as effectively changing the free energy difference ∆*ε*_AI_ between the protein’s active and inactive states (Appendix D.2), which, in turn, is akin to the role of methylation ^27,28^ or phosphorylation^28^ in adaptation, but without the covalent linkage. Our in-depth analysis of the logic repertoire available to 3-input MWC molecules can serve as a theoretical framework for designing new allosteric proteins and also for understanding the measured responses of existing systems. Examples of such systems that both act as 3-input AND gates include the GIRK channel, the state of which (open or closed) is regulated by the G protein G_β*γ*_, the lipid PIP_2_ and Na^+^ ions,^29^ or the engineered N-WASP signaling protein which is activated by SH3, Cdc42 and PDZ ligands.^30^

The exquisite control that arises from the web of interactions underlying biological systems is difficult to understand and replicate. A first step to overcoming this hurdle is to carefully quantify the types of behaviors that can arise from multi-component systems. As our ability to harness and potentially design *de novo* allosteric systems grows,^21,29–33^ we can augment our current level of combinatorial control in biological contexts, such as transcriptional regulation, ^3,14,15,34,35^ to create even richer dynamics.

## Supporting information

Jupyter Notebooks where the set of functionally unique gates and constraints conditions are derived

Mathematica note- book from which all protein activity response plots and gate quality metric plots can be reproduced

## Supporting Information Description

Details on aforementioned derivations and calculations; supplementary *Mathematica* notebook from which all protein activity response plots and gate quality metric plots can be reproduced (ZIP); supplementary Jupyter Notebooks where the set of functionally unique gates and constraints conditions are derived (ZIP).

## Acknowledgements

It is a great pleasure to acknowledge the contributions of Bill Eaton to our understanding of allostery. We thank Chandana Gopalakrishnappa and Parijat Sil for their input on this work, and Michael Elowitz for his insights and valuable feedback on the manuscript. This research was supported by La Fondation Pierre-Gilles de Gennes, the Rosen Center at Caltech, the Department of Defense through the National Defense Science & Engineering Graduate Fellowship (NDSEG) Program (LF), and the National Institutes of Health DP1 OD000217 (Director’s Pioneer Award), R01 GM085286, and 1R35 GM118043-01 (MIRA). We are grateful to the Burroughs-Wellcome Fund for its support of the Physical Biology of the Cell Course at the Marine Biological Laboratory, where part of this work was completed.

## A Derivation of Conditions for Achieving Different Logic Responses

In this section we derive the conditions necessary for an MWC molecule modulated by two ligands (with one binding site for each ligand) to exhibit the behavior of various logic gates shown in Figure 1. In addition to the three logic gates shown in Figure 1, we will also discuss the three complimentary gates NAND, NOR, and XNOR depicted in Figure S1.

**Figure S1.**
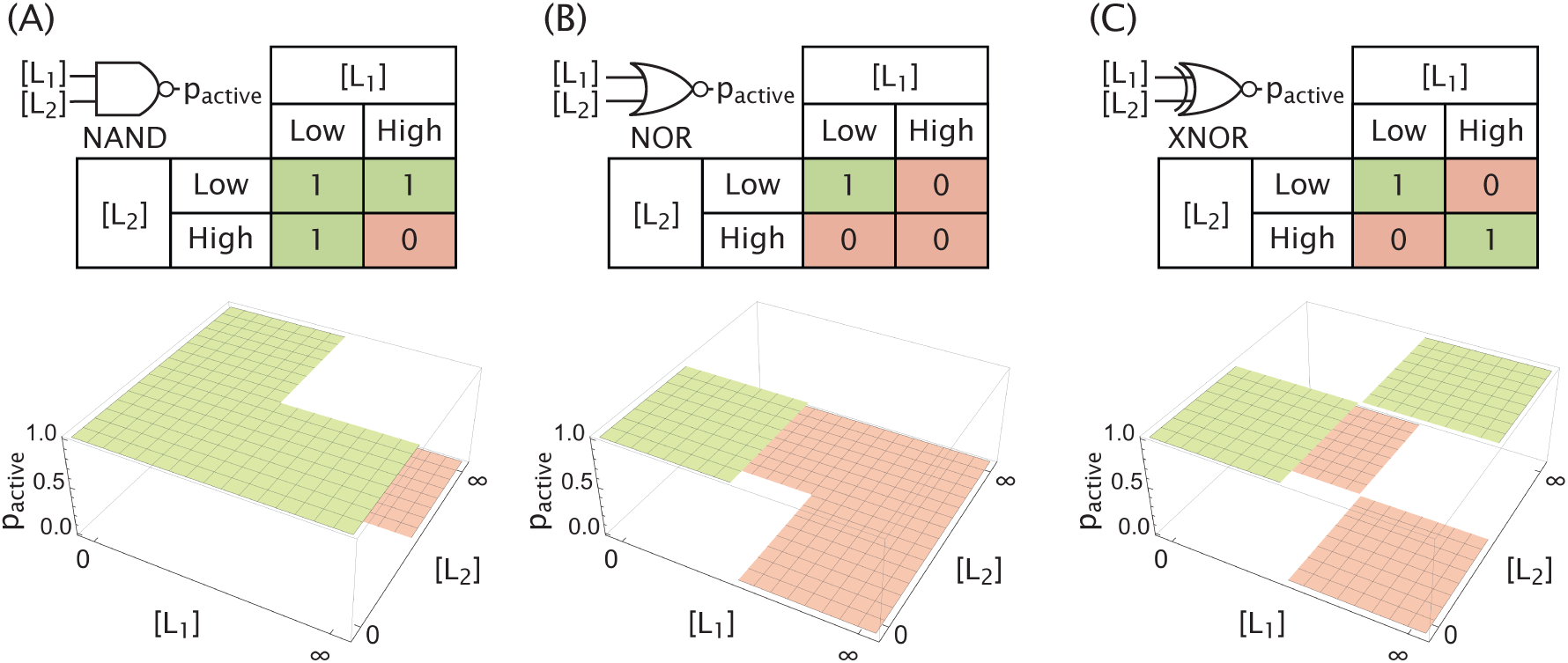
Additional logic gates as molecular responses. The (A) NAND, (B) NOR, and (C) XNOR gates are the compliments of the AND, OR, and XOR gates, respectively, shown in Figure 1.

To simplify our notation, we define the value of p_active_ from eq 1 in the following limits,

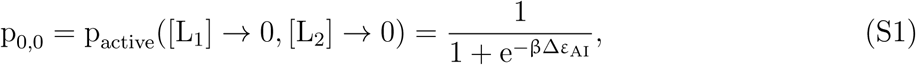

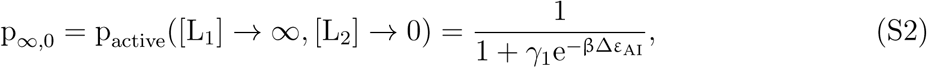

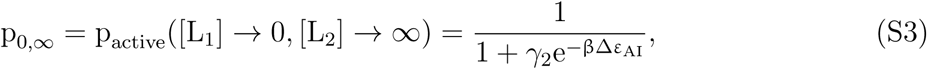

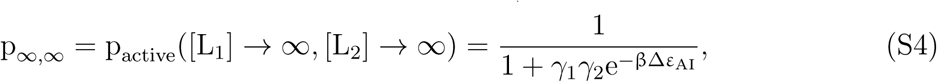

where 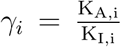 is the ratio of the dissociation constants between the i^th^ ligand and the protein in the active and inactive states. From the ideal logic gate behaviors visualized in Figure 1 and Figure S1, we can then deduce the desired constraints that model parameters need to meet for an effective realization of each gate.

## AND Gate

Starting from the AND gate, we require p_0,0_ ≈ 0, p_0,∞_ ≈ 0, p_∞,0_ ≈ 0 and p_∞,∞_ ≈ 1, which yields the following conditions:

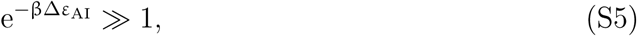

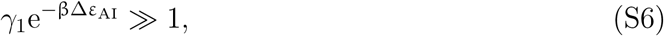

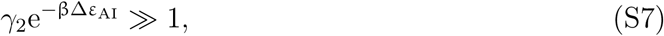

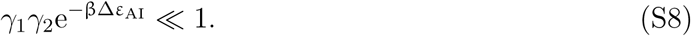

Combining eqs S6-S8, we obtain the condition for an AND gate, namely,

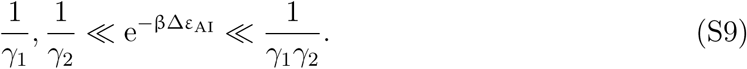

Note, that the outer inequalities imply

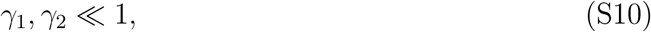

meaning that both ligands bind more tightly to the protein in the active than the inactive state.

## A.2 OR Gate

For p_active_ to represent an OR gate across ligand concentration space, it must satisfy p_0,0_ ≈ 0, p_0,∞_ ≈ 1, p_∞,0_ ≈ 1 and p_∞,∞_ ≈ 1. This requires that the parameters obey

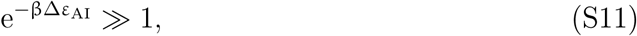

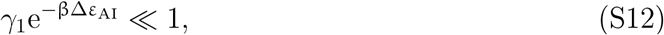

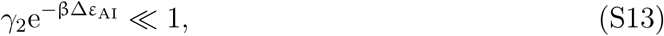

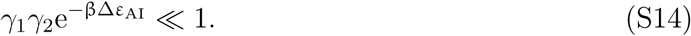

Combining eqs S11-S13, we obtain a constraint on the free energy difference,

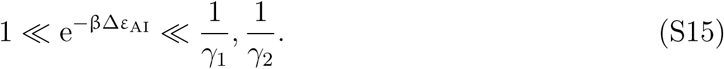

As with the AND gate, the outer inequalities imply that the ligands prefer binding to the protein in the active state,

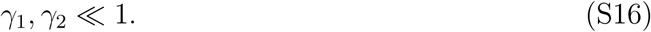

## A.3 NAND and NOR Gates

Because the NAND and NOR gates are the logical complements of AND and OR gates, respectively, the parameter constraints under which they are realized are the opposites of those for AND and OR gates. Hence, the conditions for a NAND gate are given by

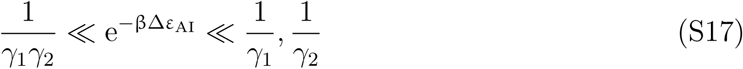

while the conditions for NOR gates are

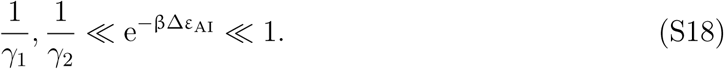

We note that in both cases, the outer inequalities imply that both ligands bind more tightly to the protein in the inactive state than in the active state, *γ*_1_, *γ*_2_ ≫ 1.

The symmetry between AND/OR and NAND/NOR gates also implies a simple relation between their quality metrics, namely, 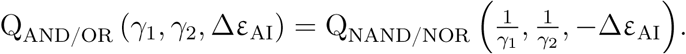.

Here we provide a proof for the AND gate and invite the reader to do the same for the OR gate. From eq 2, the quality metrics for the AND and NAND gates can be written as

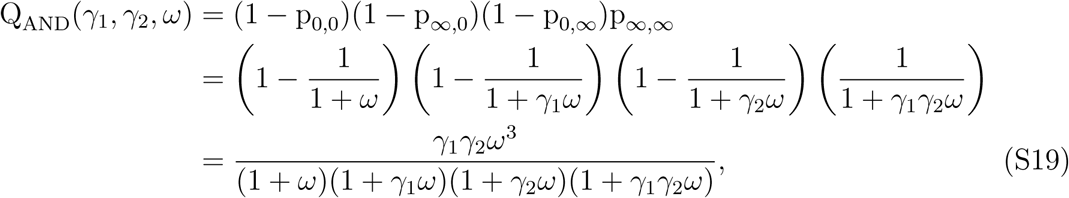

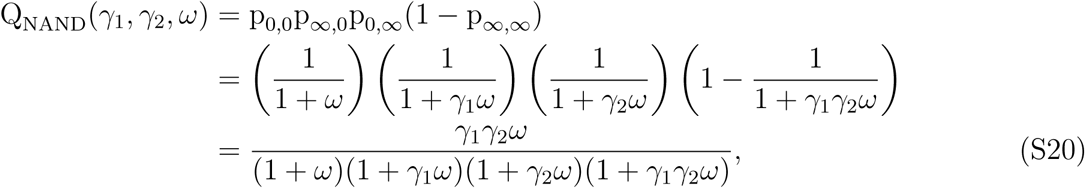

where we introduced ω = e^−β∆*ε*^^AI^. Substituting 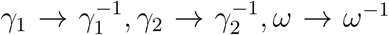 (equivalent to ∆*ε*_AI_ → *−∆*ε**_AI_) in eq S20, we obtain

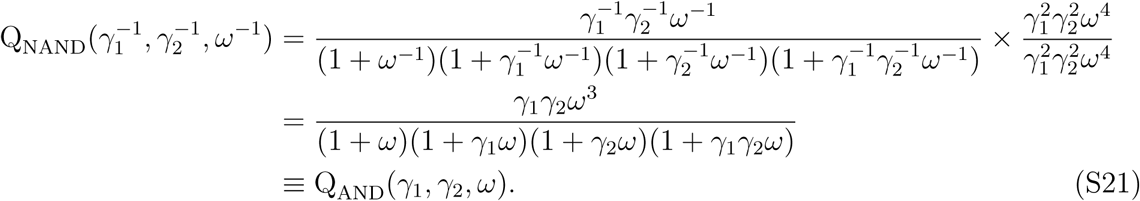

## A.4 XOR and XNOR Gates

Here, we show that the XOR gate (and by symmetry the XNOR gate) are not achievable with the form of p_active_ given in eq 1. An XOR gate satisfies p_0,0_ ≈ 0, p_0,∞_ ≈ 1, p_∞,0_ ≈ 1 and p_∞,∞_ ≈ 0 which necessitates the parameter conditions

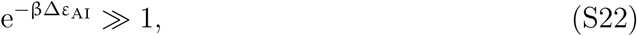

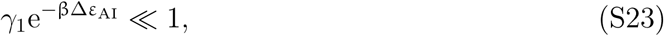

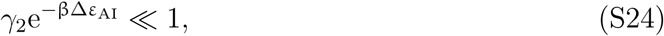

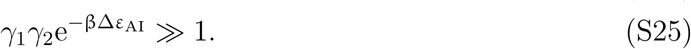

However, these conditions cannot all be satisfied, as the left-hand side of eq S25 can be written in terms of the left-hand sides of eqs S22-S24,

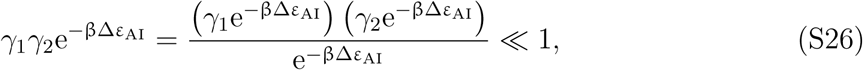

contradicting eq S25.

The XOR gate could be realized if an explicit cooperativity energy *ε*_A,coop_ is added when both ligands are bound in the active state and *ε*_I,coop_ when both are bound in the inactive state. These cooperative interactions modify eq 1 to the form

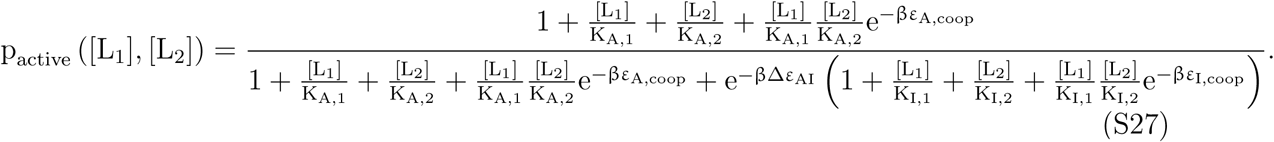

Figure S2 demonstrates that the same parameter values from Figure 3B together with the (unfavorable) cooperativity energy *ε*_A,coop_ = 15 k_B_T and *ε*_I,coop_ = 0 can create an XOR gate.

**Figure S2.**
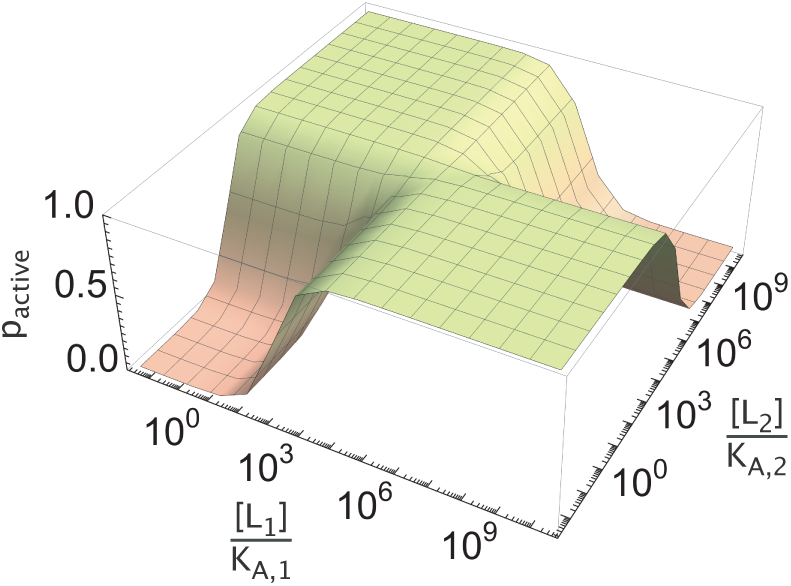
An XOR gate can be achieved by adding cooperativity. The activity profile defined in eq S27 for the parameter values from Figure 3B, along with the cooperativity energies *ε*_A,coop_ = 15 k_B_T and *ε*_I,coop_ = 0, give rise to an XOR response.

## B The General Two-Ligand Response: Transitioning Between OFF and ON States

In the preceding section, we have been solely concerned with the behavior of the MWC molecule in the limits of ligand concentration ([L_i_] = 0 and [L_i_] → ∞), and have ignored the details about the transition from ON to OFF (e.g., its shape and steepness) and also the possibility of p_active_ ≠ 0 or 1. In this section, we examine and derive in greater detail some of the additional response behaviors that are possible for an MWC molecule regulated with N = 2 ligands when the locations of transitions between limit responses are taken into account.

To examine the transitions between p_active_ levels, we derive expressions for the concentrations at which transitions are at their midpoint. Since p_active_ is a function of two different ligand concentrations, [L_1_] and [L_2_], we define two different midpoint concentrations of ligand L_i_: one in the absence of ligand L_j_, 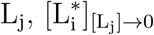, and another when L_j_ is saturating, 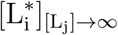. In particular, 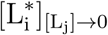 is defined such that

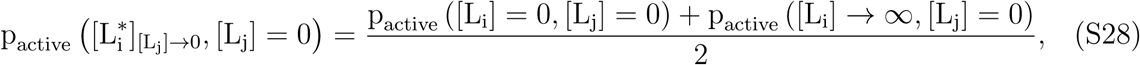

i.e., the concentration of ligand i where p_active_ is equal to the mean of the two p_active_ limit values being transitioned between. If we evaluate the left hand side of eq S28 with i = 1 and j = 2 using eq 1, and the right hand side using the limits from Figure 3(A), we obtain

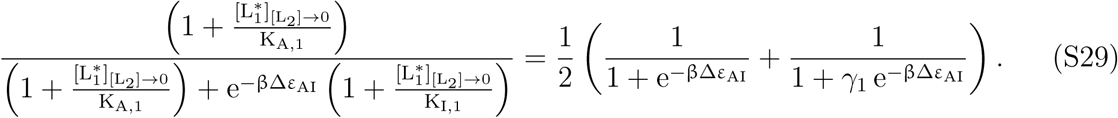

Introducing *γ*_1_ = K_A,1_*/*K_I,1_, we can solve for 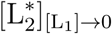 to find

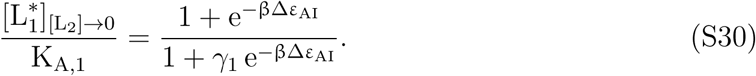

Eq S30 can be rewritten for [L∗_2_]_[L1]→0_ by merely interchanging all ligand and parameter indices, i.e., 1 ↔ 2.

The midpoint concentration when one ligand is saturating can be derived similarly. Specifically, to find an expression for 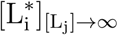 we can re-write S28 using eq 1 in the case that [L_j_] → ∞ with i = 1 and j = 2, resulting in

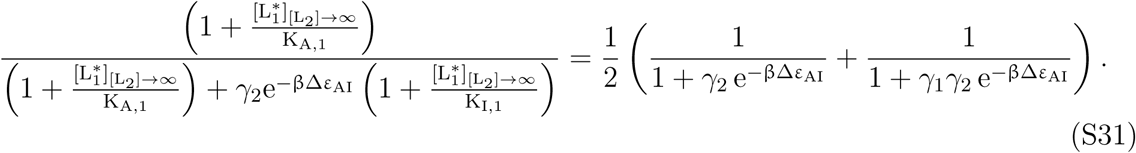

Eq S31 can be solved for 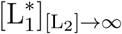 to produce,

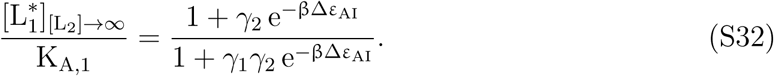

Again, the symmetric expression for 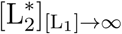 is found by swapping ligand and parameter indices, 1 ↔ 2.

Using this approach to define concentration transition zones can be used to produce additional MWC behaviors, including the ratiometric response in the BMP pathway recently analyzed by Antebi *et al.*,^8^ which was briefly discussed earlier. Specifically, this response can be approximated by choosing parameter values that satisfy two desired limits, p∞_,0_ ≈ 0 (*γ*_1_ e−β∆*ε*^AI »^ 1) and p_0,∞_ ≈ 1 (*γ*_2_ e−β∆*ε*^AI «^ 1), as well as produce a large transition region sensitive to both ligands, i.e., the ratio in eq 7, 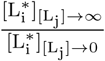 is far from 1. One way to satisfy these conditions is to set K_I,2_ K_A,1_ = K_A,2_ K_I,1_ and ∆*ε*_AI_ = 0 in eq 1. Notice that with these parameter choices and provided the ligand concentrations satisfy

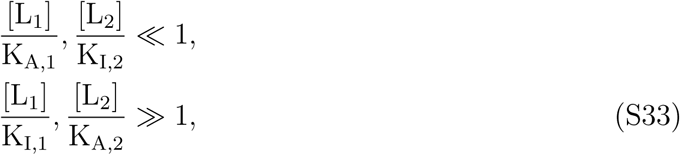

the probability that the protein is active reduces to

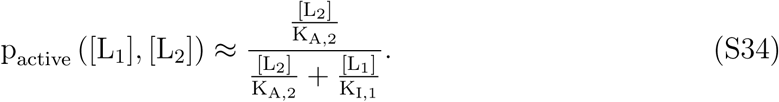

Hence, only the ratio of [L_1_] and [L_2_] matters, as shown in Figure 4B where eq S33 is satisfied provided that 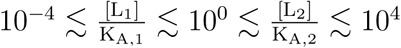

Additionally, we consider the remaining three types of input-output computations shown by Antebi *et al.* to exist in the BMP pathway which they called the additive, imbalance, and balance responses.^8^ The additive response (which responds more to larger input concentrations) is an OR gate which we showed is possible in Figure 3B. The imbalance response (which responds maximally to extreme ratios of the two input ligands) is similar to an XOR behavior which, as discussed in Appendix A.4, is only achievable with an explicit cooperativity energy.

The balance response is defined as

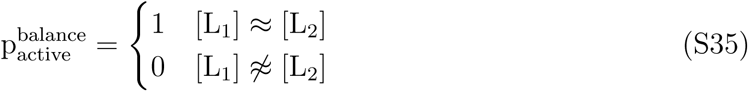

so that the protein is only ON when both ligands are present in the same amount as shown in Figure S3A. Such behavior is not possible within the MWC model because starting from any point 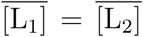, p_active_ in eq 1 must either monotonically increase or monotonically decrease with [L_1_] (depending on *γ*_1_), whereas eq S35 requires that p_active_ must decrease for both 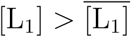 and 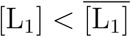 (with similar contradictory statements for [L_2_]). The closest behavior achievable by the MWC model is to zoom into the transition region of an XNOR gate as shown in Figure S3B. As we zoom out of the concentration ranges shown, the four square regions of the plot will continue to expand as squares and the behavior will no longer approximate the ideal balance response.

**Figure S3.**
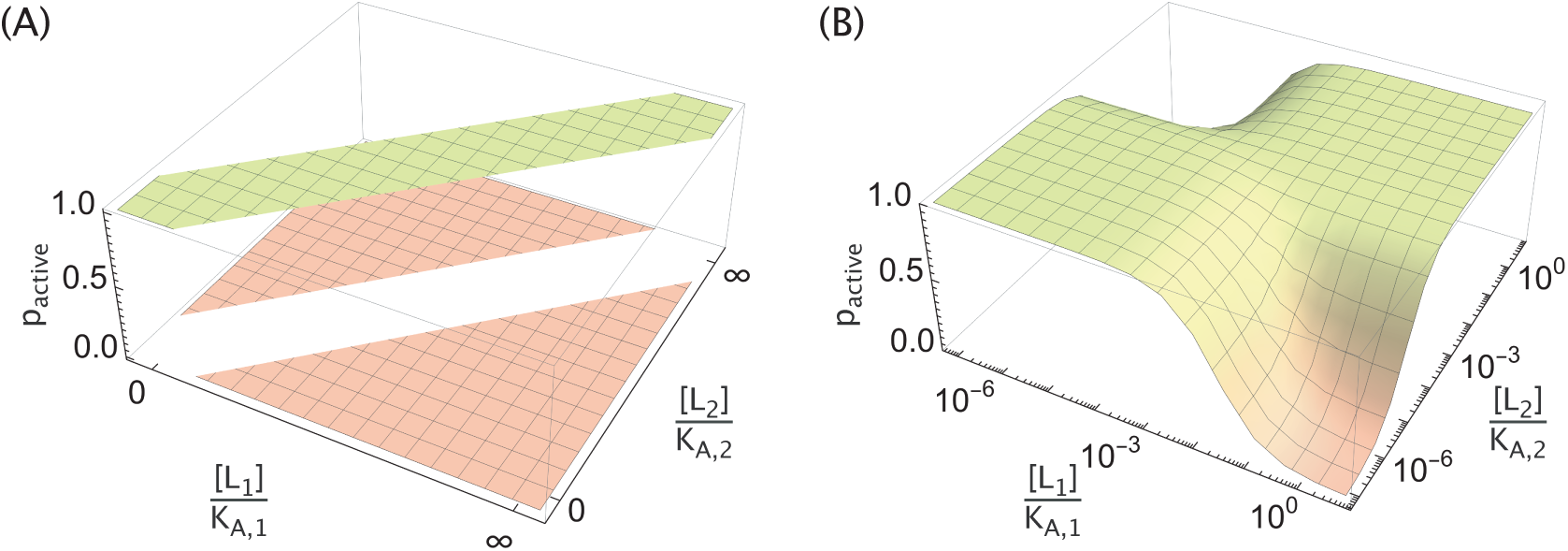
Balance response behavior approximated by the MWC model. (A) The ideal balance response from the BMP pathway and (B) the closest behavior that an MWC molecule can exhibit using the complementary parameters from Figure S2 (K_A,i_ = 1.5 × 10^−4^ M, K_I,i_ = 2.5 × 10^−8^ M, ∆*ε*_AI_ = 5 k_B_T, *ε*_A,coop_ = −15 k_B_T and *ε*_I,coop_ = 0).

## Logic Switching by Tuning the Number of Ligand Binding Sites

In this section, we show how an MWC molecule whose activity is given by eq 9 can switch between exhibiting AND↔OR or NAND↔NOR behaviors by tuning the number of binding sites. To begin, we define the probability p_active_ that the molecule is active in the case when the i^th^ ligand has n_i_ binding sites, namely,

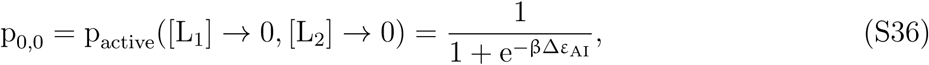

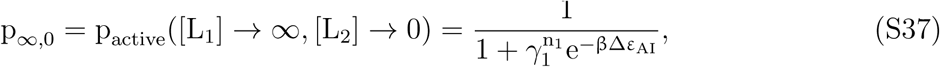

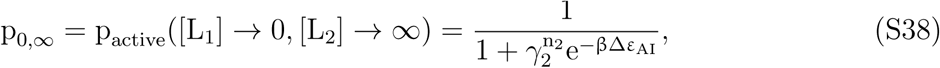

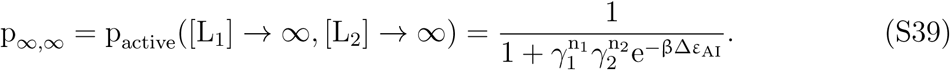

Note that the only effect of having an arbitrary number of ligand binding sites (as opposed to n_i_ = 1 as in Appendix A) is that the ratio of dissociation constants always appears raised to the number of binding sites, 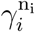. Hence, the parameter conditions derived for AND and OR behaviors for n_i_ = 1 can be used in the case of general n_i_ by substituting 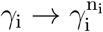

Now, suppose a molecule with N = 2 ligands and with 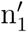 and 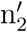 binding sites for ligands 1 and 2 represents an AND gate, while this same molecule with n_1_ and n_2_ binding sites serves as an OR gate, as in Figure 5B with 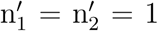 and n_1_ = n_2_ = 4. From Figure 3B, the conditions in the former case (AND gate) are

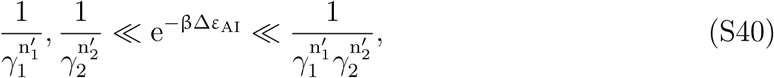

while the conditions in the latter case (OR gate) are

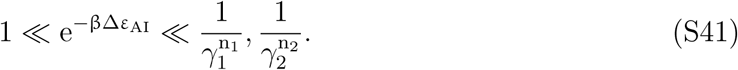

Combining these conditions, we find that the requirements for the AND ↔ OR switching are given by

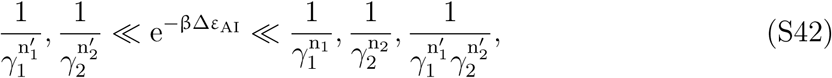

where we have used the fact that the outer inequalities imply 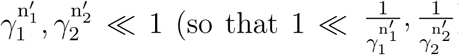. In the limit 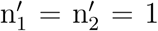, eq S42 reduces to the condition shown in Figure 5A.

Lastly, we note that since NAND is the complement of AND while NOR is the complement of OR, the class switching requirements in S42 become the requirements to change from NAND↔NOR behavior when 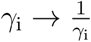 and ∆*ε*_AI_ → *−∆*ε**_AI_.

## D Combinatorial Control with Three Regulatory Ligands

In this section, we first present the methodology used to identify the functionally unique and MWC-compatible 3-ligand logic gates. We then use the full list of admissible gates to find all possible logic switches that can be induced by increasing the concentration of a third ligand. We finish the section by deriving the parameter conditions required for achieving the logic switches AND→OR and AND→YES_1_ shown in Figure 7D.

### D.1 Functionally Unique MWC Gates

To identify the set of functionally unique MWC gates, we first iterate over the 256 possible responses and eliminate those redundant ones that can be obtained by shuffling the ligand labels of already selected gates. The Python implementation of this procedure that leaves 80 functionally unique gates can be found in the supplementary Jupyter Notebook 1.

Having singled out the functionally unique responses, we proceed to identify those that are admissible in the MWC framework. To that end, we first write the analytic forms for the probability of the protein being active (p_active_) at eight different ligand concentration limits (Figure S4A). Since the functional form in all cases is p_active_ = (1 + w_I/A_)^−1^, where w_I/A_ is the total weight of the inactive states divided by the total weight of the active states in the appropriate limit (as seen in Figure 3A), a Boolean response (p_active_ ≈ 0 or 1) can only be achieved when w_I/A_ ≫ 1 or w_I/A_ ≪ 1, respectively. Hence, the values of w_I/A_ at the eight different limits of ligand concentration will determine the full logic response of the protein.

Note that since cooperative interactions between ligands are absent in the MWC framework, the eight different w_I/A_ expressions depend on only four independent MWC parameters, namely,{∆*ε*_AI_, *γ*_1_, *γ*_2_, *γ*_3_}. Therefore, only four of the eight limiting w_I/A_ values can be independently tuned, and any w_I/A_ limit can be expressed as a function of four different and independent w_I/A_ limits, resulting in a constraint condition. Since each w_I/A_ is a product of some *γ*_i_’s and e^−β∆*ε*^_AI_ (Figure S4A), we look for constraint conditions that have a multiplicative form, namely,

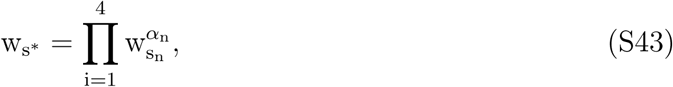

where w_s_* is the target limit, s_n_ ≠ s∗(1 ≤ n ≤ 4) are the labels of four different limits and *α*_n_ are real coefficients. Searching over all conditions of such form (see the supplementary Jupyter Notebook 2 for details), we identify a total of eight functionally unique constraints,

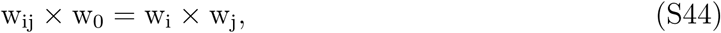

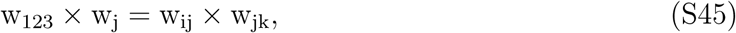

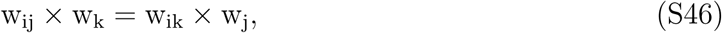

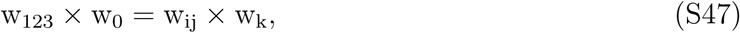

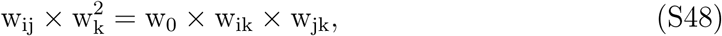

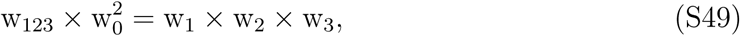

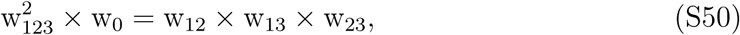

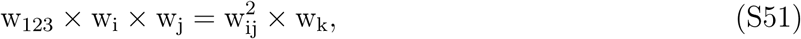

where 1 ≤ i, j, k ≤ 3.

Further searching for a minimum set of constraints that can account for all gates incompatible with the MWC framework, we identify the constraints in eqs S44-S47 as the necessary and sufficient ones (see the supplementary Jupyter Notebook 2). Graphical representations of these four constraints on a cubic diagram are shown in Figure S4B. Note that these conditions are all of the form

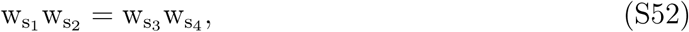

where s_i_ are labels corresponding to different ligand concentration limits. Logic responses where w_s1_, w_s2_ ≪ 1 (≫ 1) while w_s3_, w_s4_ ≫ 1 (≪ 1) cannot be achieved, since they contradict the constraint condition. Conditions 1 and 2 in Figure S4B, for example, demonstrate that XOR and XNOR gates cannot be realized by any two ligands in the absence (condition 1) or presence (condition 2) of a third ligand - a result expected from the 2-ligand analysis done earlier. On the other hand, conditions 3 and 4 are specific to the 3-ligand response.

Checking the 80 functionally unique gates against the four constraints in Figure S4B, we obtain a set of 34 functionally unique and MWC-compatible gates, 17 of which are shown in Figure S5A while the other half are their logical complements (i.e. ON↔OFF swapping is performed for each of the cube elements).

**Figure S4.**
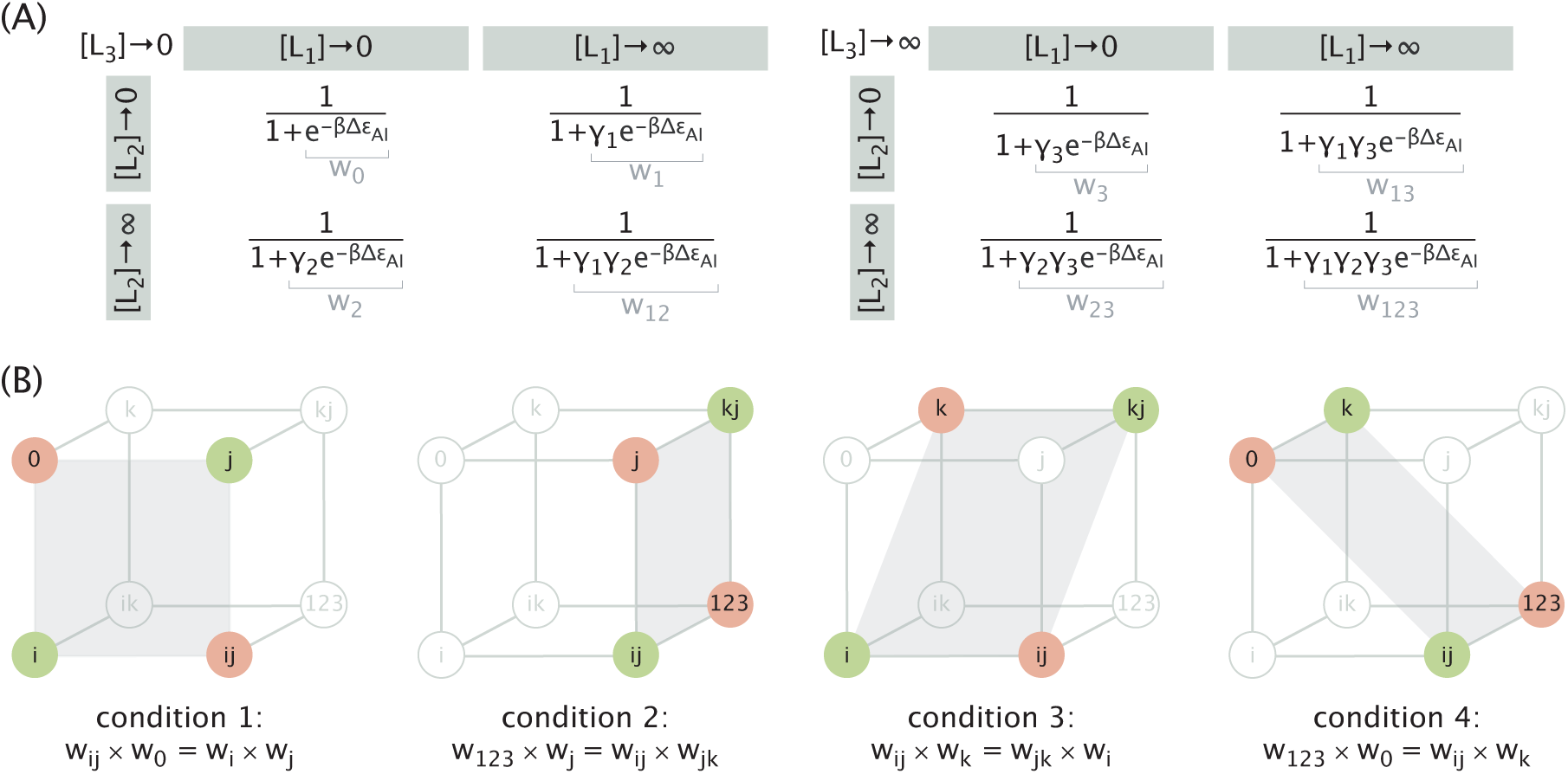
Three-ligand logic gates that are incompatible with the MWC framework. (A) Probability that the protein is active in the 8 different ligand concentration limits. The total weight of the inactive states relative to the active states is indicated in gray for all limits. (B) Cubic diagrams of logic responses that are incompatible with the MWC framework, along with the constraint equations used to obtain them. The limits relevant to the constraint conditions are shown in color, and a transparent gray plane containing these relevant limits is added for clarity. In all four diagrams 1 ≤ i, j, k ≤ 3.

## D.2 Logic Switching

Here we describe how the table of all possible logic switches inducible by a third ligand (Figure 6D) can be obtained from the list of MWC-compatible 3-ligand gates (Figure S5), and also derive the parameter conditions for AND→OR and AND→YES_1_ logic switches.

As illustrated in Figure 6C, logic switching can be achieved by increasing the concentration of any of the three ligands. Following the same procedure, we iterate over the list of gates shown in Figure S5A and for each of them identify the set of possible logic switches. The set of all logic switches present in Figure S5A together constitute the entries of the table in Figure 6D. Note that if a gate is compatible with the MWC framework, then its logical complement is also compatible, and therefore, the possibility of switching between two gates, Gate 1 → Gate 2, implies the possibility of switching between their logical complements, NOT → (Gate 1) → NOT → (Gate 2).

**Figure S5.**
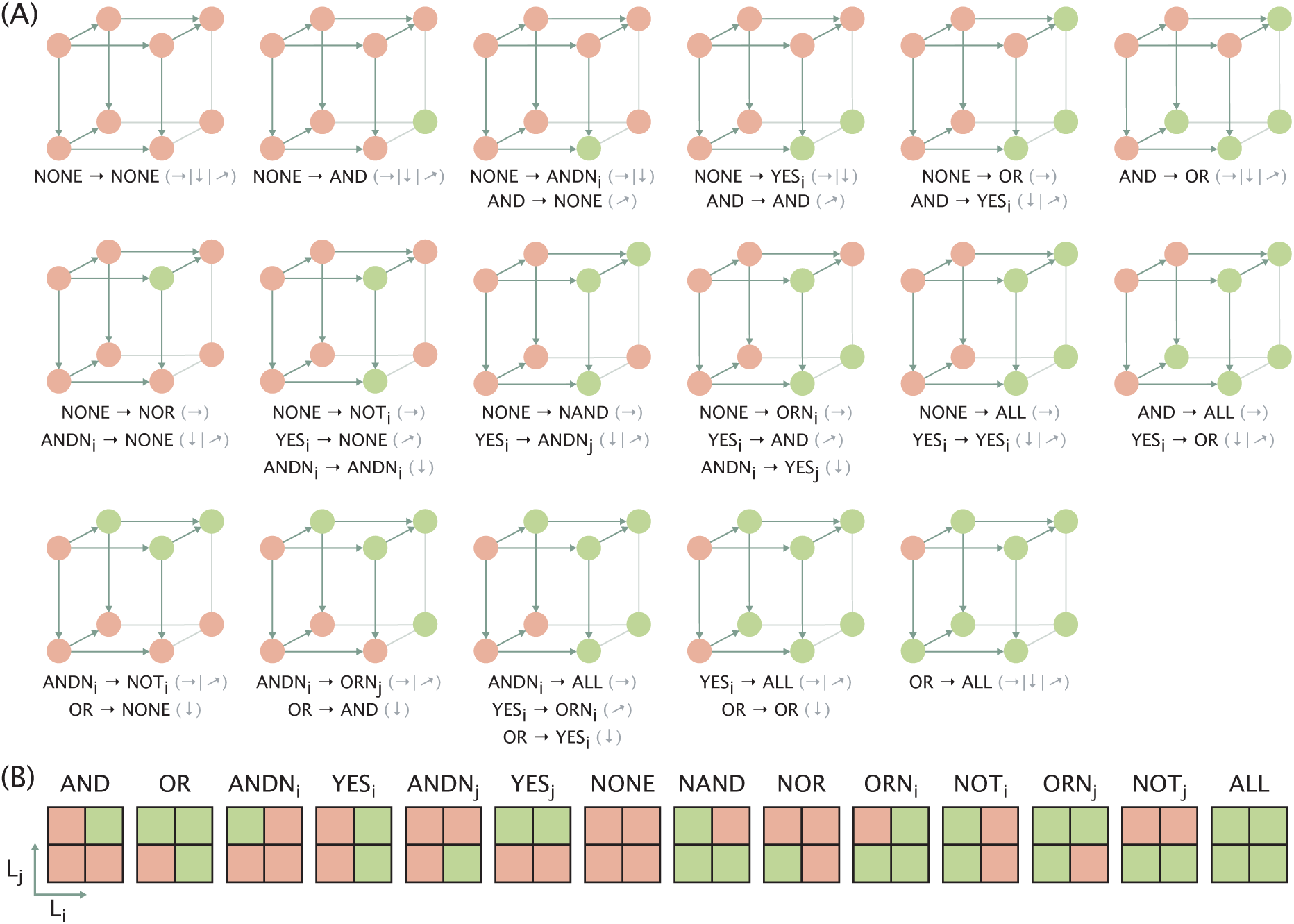
Functionally unique 3-ligand MWC gates and possible schemes of logic switching. (A) List of functionally unique 3-ligand MWC gates that have an inactive base state (in the absence of ligands). The set of logic switches that can be achieved by increasing the concentration of one of the ligands is listed on the bottom of each gate, with the gray arrows indicating the corresponding directions of increasing ligand concentration. Transitions with swapped labels (i ↔ j) are also possible and are not listed. Arrows corresponding to the ligand axes on different faces of the cube are included to assist the derivation of possible logic switches. (B) Schematics of 2-ligand gates adapted from Figure 6D for convenience.

Now, we show how an MWC protein can exhibit the switching behaviors in Figure 7B,D (AND→OR and AND→YES_1_) by saturating the concentration of the third ligand. We first consider the behavior of the protein in the absence of the third ligand ([L_3_] = 0, with p_active_ limits given in Figure S4A, left) and then consider how the protein acts at the saturating concentration of the third ligand ([L_3_] → ∞, with p_active_ limits given in Figure S4A, right). With [L_3_] = 0, the protein ignores the third ligand and behaves identically to a protein with N = 2 ligands. In the limit [L_3_] → *∞*, however, the protein behaves as if it only has two ligands with an altered free energy difference 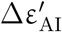 between the active and inactive states given by

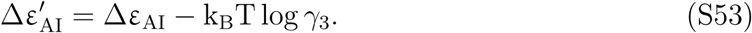

Suppose that a protein acts as an AND gate when [L_3_] = 0 and transitions into an OR gate when [L_3_] → ∞, as in Figure 7B. From Figure 3B, the MWC parameters must satisfy

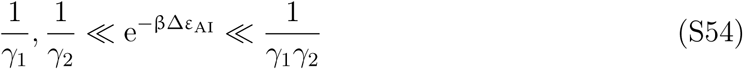

in the absence of L_3_ (AND behavior) and

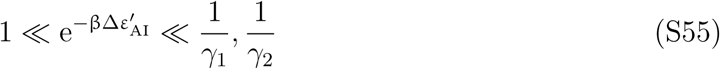

when [L_3_] is saturating (OR behavior). Using eqs S53, we can rewrite the condition S55 as

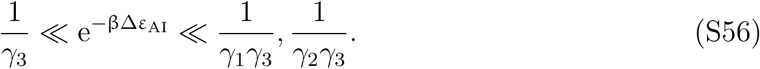

Combining eq S54 and eq S56, we find the second condition reported in Figure 7A, namely,

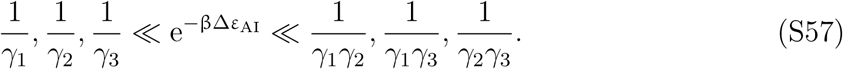

The first condition in Figure 7A is then obtained by using the outer inequalities, that is,

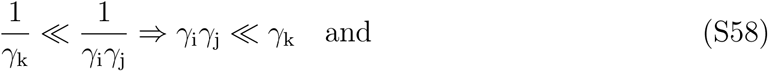

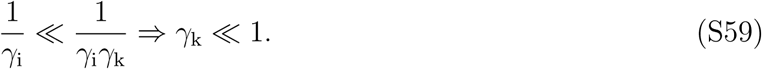

Lastly, we derive the parameter conditions needed to achieve an AND→YES_1_ switching by saturating the third ligand. Conditions for the AND behavior in the absence of the third ligand are already known (eq S54). To achieve a YES_1_ gate, p_active_ at [L_3_] → ∞ needs to meet the following limits:

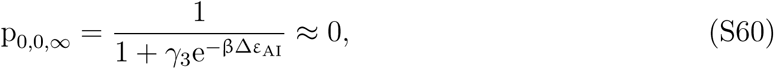

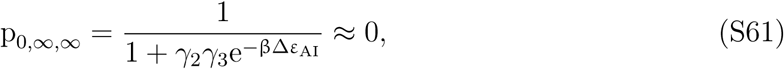

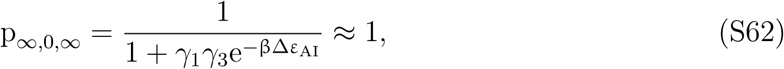

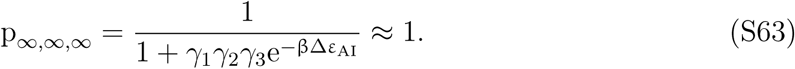

These limits suggest constraints on ∆*ε*_AI_, which, combined with eq S54, result in

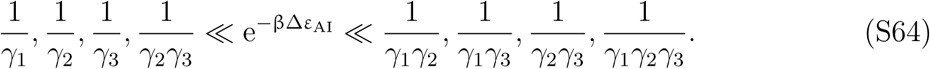

The outer inequalities, in turn, suggest conditions for the *γ* parameters, namely,

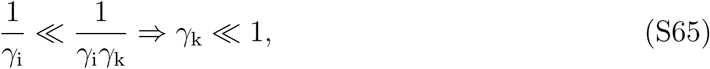

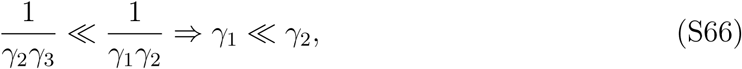

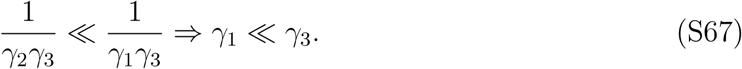

Accounting for these additional constraints, eq S64 simplifies into

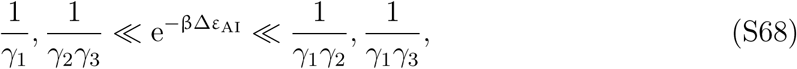

as shown in Figure 7C.

## References

(1) Blangy, D.; Buc, H.; Monod, J. Kinetics of the Allosteric Interactions of Phosphofructokinase from Escherichia Coli. Journal of Molecular Biology 1968, 31, 13–35.

(2) Dueber, J. E.; Yeh, B. J.; Chak, K.; Lim, W. A. Reprogramming Control of an Allosteric Signaling Switch Through Modular Recombination. Science 2003, 301, 1904–1908.

(3) Buchler, N. E.; Gerland, U.; Hwa, T. On Schemes of Combinatorial Transcription Logic. Proceedings of the National Academy of Sciences 2003, 100, 5136–41.

(4) Graham, I.; Duke, T. The Logical Repertoire of Ligand-Binding Proteins. Physical Biology 2005, 2, 159–165.

(5) de Ronde, W.; ten Wolde, P. R.; Mugler, A. Protein Logic: A Statistical Mechanical Study of Signal Integration at the Single-Molecule Level. Biophysical Journal 2012, 103, 1097–1107.

(6) Agliari, E.; Altavilla, M.; Barra, A.; Schiavo, L. D.; Katz, E. Notes on Stochastic (Bio)-Logic Gates: Computing With Allosteric Cooperativity. Scientific Reports 2015, 5, 9415.

(7) Martins, B. M.; Swain, P. S. Trade-offs and Constraints in Allosteric Sensing. PLoS Computational Biology 2011, 7, e1002261.

(8) Antebi, Y. E.; Linton, J. M.; Klumpe, H.; Bintu, B.; Gong, M.; Su, C.; McCardell, R.; Elowitz, M. B. Combinatorial Signal Perception in the BMP Pathway. Cell 2017, 170, 1184–1196.

(9) Escalante-Chong, R.; Savir, Y.; Carroll, S. M.; Ingraham, J. B.; Wang, J.; Marx, C. J.; Springer, M. Galactose Metabolic Genes in Yeast Respond to a Ratio of Galactose and Glucose. Proceedings of the National Academy of Sciences 2015, 112, 1636–1641.

(10) Razo-Mejia, M.; Barnes, S. L.; Belliveau, N. M.; Chure, G.; Einav, T.; Lewis, M.; Phillips, R. Tuning Transcriptional Regulation through Signaling: A Predictive Theory of Allosteric Induction. Cell Systems 2018, 6, 456–469.

(11) Auerbach, A. Thinking in Cycles: MWC Is a Good Model for Acetylcholine Receptor-Channels. The Journal of Physiology 2012, 590, 93–98.

(12) Marzen, S.; Garcia, H. G.; Phillips, R. Statistical Mechanics of Monod-Wyman-Changeux (MWC) Models. Journal of Molecular Biology 2013, 425, 1433–1460.

(13) Mirny, L. A. Nucleosome-Mediated Cooperativity Between Transcription Factors. Proceedings of the National Academy of Sciences 2010, 107, 22534–22539.

(14) Scholes, C.; DePace, A. H.; Sánchez, Á. Combinatorial Gene Regulation through Kinetic Control of the Transcription Cycle. Cell Systems 2017, 4, 97–108.

(15) Kinkhabwala, A.; Guet, C. C. Uncovering Cis Regulatory Codes Using Synthetic Promoter Shuffling. PLoS ONE 2008, 3, e2030.

(16) Dueber, J. E.; Yeh, B. J.; Bhattacharyya, R. P.; Lim, W. A. Rewiring Cell Signaling: The Logic and Plasticity of Eukaryotic Protein Circuitry. Current Opinion in Structural Biology 2004, 14, 690–699.

(17) Privman, V.; Zhou, J.; Halámek, J.; Katz, E. Realization and Properties of Biochemical-Computing Biocatalytic XOR Gate Based on Signal Change. The Journal of Physical Chemistry B 2010, 114, 13601–13608.

(18) de Silva, A. P.; McClenaghan, N. D. Simultaneously Multiply-Configurable or Superposed Molecular Logic Systems Composed of ICT (Internal Charge Transfer) Chromophores and Fluorophores Integrated with One- or Two-Ion Receptors. Chemistry-A European Journal 2002, 8, 4935–4945.

(19) De Silva, A. P.; Uchiyama, S. Molecular Logic and Computing. Nature Nanotechnology 2007, 2, 399.

(20) Bloom, J. D.; Meyer, M. M.; Meinhold, P.; Otey, C. R.; MacMillan, D.; Arnold, F. H. Evolving Strategies for Enzyme Engineering. Current Opinion in Structural Biology 2005, 15, 447–452.

(21) Guntas, G.; Ostermeier, M. Creation of an Allosteric Enzyme by Domain Insertion. Journal of Molecular Biology 2004, 336, 263–273.

(22) Narula, J.; Igoshin, O. A. Thermodynamic Models of Combinatorial Gene Regulation by Distant Enhancers. IET Systems Biology 2010, 4, 393–408.

(23) Löhr, U.; Chung, H.-R.; Beller, M.; Jäckle, H. Antagonistic Action of Bicoid and the Repressor Capicua Determines the Spatial Limits of Drosophila Head Gene Expression Domains. Proceedings of the National Academy of Sciences 2009, 106, 21695–21700.

(24) Fakhouri, W. D.; Ay, A.; Sayal, R.; Dresch, J.; Dayringer, E.; Arnosti, D. N. Deciphering a Transcriptional Regulatory Code: Modeling Short-Range Repression in the Drosophila Embryo. Molecular systems biology 2010, 6, 341.

(25) Chen, H.; Xu, Z.; Mei, C.; Yu, D.; Small, S. A System of Repressor Gradients Spatially Organizes the Boundaries of Bicoid-Dependent Target Genes. Cell 2012, 149, 618–629.

(26) Crocker, J.; Tsai, A.; Stern, D. L. A Fully Synthetic Transcriptional Platform for a Multicellular Eukaryote. Cell Reports 2017, 18, 287–296.

(27) Hansen, C. H.; Endres, R. G.; Wingreen, N. S. Chemotaxis in Escherichia Coli: A Molecular Model for Robust Precise Adaptation. PLoS Computational Biology 2008, 4, e1.

(28) Lan, G.; Sartori, P.; Neumann, S.; Sourjik, V.; Tu, Y. The Energy-Speed-Accuracy Trade-Off in Sensory Adaptation. Nature Physics 2012, 8, 422.

(29) Wang, W.; Touhara, K. K.; Weir, K.; Bean, B. P.; MacKinnon, R. Cooperative Regulation by G Proteins and Na(+) of Neuronal GIRK2 K(+) Channels. eLife 2016, 5, e157519.

(30) Dueber, J. E.; Mirsky, E. A.; Lim, W. A. Engineering Synthetic Signaling Proteins with Ultrasensitive Input/Output Control. Nature Biotechnology 2007, 25, 660–662.

(31) Raman, A. S.; White, K. I.; Ranganathan, R. Origins of Allostery and Evolvability in Proteins: A Case Study. Cell 2016, 166, 468–480.

(32) Huang, P. S.; Boyken, S. E.; Baker, D. The Coming of Age of De Novo Protein Design. Nature 2016, 537, 320.

(33) Guntas, G.; Mansell, T. J.; Kim, J. R.; Ostermeier, M. Directed Evolution of Protein Switches and Their Application to the Creation of Ligand-Binding Proteins. Proceedings of the National Academy of Sciences 2005, 102, 11224–11229.

(34) Wei, H.; Hu, B.; Tang, S.; Zhao, G.; Guan, Y. Repressor Logic Modules Assembled by Rolling Circle Amplification Platform to Construct a Set of Logic Gates. Scientific Reports 2016, 6, 37477.

(35) Macía, J.; Posas, F.; Solé, R. Distributed Computation: The New Wave of Synthetic Biology Devices. Trends in Biotechnology 2012, 30, 342–349.

