## Supplementary material for "Combinatorial Control through Allostery": Jupyter Notebooks where the set of functionally unique gates and constraints conditions are derived: functionally_unique_3input_gates.ipynb

```
# %% [markdown]
In this notebook, we derive the set of functionally unique 3-input gates using the "minterm" approach. 

# %% [code]
from sympy.logic import *
from sympy import symbols
import pandas as pd
import itertools
from tqdm import tqdm_notebook as tqdn

# %% [markdown]
### Create list of all possible inputs

The possible combinations of "inputs" to the system are represented by each row of a logic table. Thus, for an n = 3 input system, there are $2^n$ = $2^3$ = 8 possible combinations of inputs:

| Input 1 | Input 2 | Input 3 |
|---------|---------|---------|
|    0    |    0    |    0    |
|    1    |    0    |    0    |
|    1    |    1    |    0    |
|    1    |    0    |    1    |
|    0    |    1    |    0    |
|    0    |    1    |    1    |
|    0    |    0    |    1    |
|    1    |    1    |    1    |


# %% [code]
input_combinations = [[input1,input2,input3] for input1 in [0,1] for input2 in [0,1] for input3 in [0,1]]
print('Total input combinations produced by code = '+ str(len(input_combinations)))

# %% [markdown]
### Create list of all possible 3 input logic responses

One way to represent a logic response or logic gates is by using "minterms," an approach also known as canonical disjunctive normal form (CDNF). This method characterizes a logical response by a list of the input combinations that produces a "1" output (as opposed to a "0" output). This allows easy conversion of logic tables into a form that lends itself well to comparing logic responses. As an example, the logic response represented by the logic table,

| Input 1 | Input 2 | Input 3 |  Output |
|---------|---------|---------|---------|
|    0    |    0    |    0    |  **0**  |
|    1    |    0    |    0    |  **0**  |
|    1    |    1    |    0    |  **1**  |
|    1    |    0    |    1    |  **1**  |
|    0    |    1    |    0    |  **0**  |
|    0    |    1    |    1    |  **1**  |
|    0    |    0    |    1    |  **0**  |
|    1    |    1    |    1    |  **1**  |

can be succinctly written as a list of its minterms:[[1,1,0], [1,0,1], [0,1,1], [1,1,1]]. Note that the list of all possible minterms is the same as the list of all possible input combinations produced earlier.

Thus creating a list of all possible 3 input logic responses is equivalent to creating a list of all possible combinations of minterms (the list of minterms can be of any length: 0 minterms $\to$ 8 minterms). Since there are m = 8 total minterms and each minterm can be either included or excluded (2 output states possible) when representing a logic response, we expect there to be $2^m$ = $2^8$ = 256 total combinations of minterms (total possible 3 input logic responses).

# %% [code]
#create an empty list to store combinations of minterms produced in the for loop 
minterm_combinations = []

#include the no minterm case (where there are no "on", i.e., "1" output responses)
minterm_combinations.append([])

#iterate over all possible lengths of minterm combinations
for combination_length in range(1,len(input_combinations)+1):
    
    #for given length of minterm combinations, list and append all possible combinations of minterms
    for combination in list(itertools.combinations(input_combinations,combination_length)):
        minterm_combinations.append(list(combination))

print('Total minterm combinations (3 input logic responses) produced by code = '+ str(len(minterm_combinations)))

# %% [markdown]
# Reduce list of all possible 3 input responses to those that are "functionally unique"

We define functional uniqueness between two logic responses if rearranging the input variable names of one logic response cannot reproduce the other logic function. As an example, the 3 input logic responses "Input_1 AND Input_2" and "Input_2 AND Input_3" are considered not unique, since replacing ("mapping") the input variable name "Input_1" in the first logic response with "Input_3" reproduces the second logic response. The logic tables of these responses can be examined for further clarity on this subject: rearranging the header labels of the input columns is equivalent to mapping input variables.

To evaluate the functional uniqueness of our generated list of minterm combinations, we convert minterm combinations to Boolean logic expressions, then write a function to compare two logic expressions. This function is used to do pairwise comparison of all logic functions that have the same number of minterms. Two logic expressions that have different numbers of minterms are by definition not mappable.

Note that there is a sympy.logic function, boolalg.bool_map(), that does something similar to our custom logic comparison function but has some differences in the definition of uniqueness, in particular negation of a variable is allowed when mapping between logic expressions, and thus cannot be used for our purposes.

Also note: all further scripting will be completed using dataframes from the pandas package for organizational purposes. 

# %% [code]
# make list corresponding to minterm_combinations that reports whether a minterm combination contains 
# the [0,0,0] minterm. This is for organizational purposes later on.
zeros_minterm = [[0,0,0] in minterm_list for minterm_list in minterm_combinations]

#convert list of all possible minterm combinations into a pandas dataframe
df = pd.DataFrame({'minterms':minterm_combinations,'zeros_minterm_present':zeros_minterm})
df

# %% [code]
# create symbolic variables L_1, L_2, L_3 representing the three input ligands
L_1,L_2,L_3 = symbols('L_1 L_2 L_3')
variable_list = [L_1,L_2,L_3]

# add a column to our dataframe for the number of minterms for each logic function
df['num_minterms']=df.apply(lambda row: len(row.minterms),axis=1)

#rearrange columns
df = df[['num_minterms','minterms','zeros_minterm_present']]
df.head(30)

# %% [markdown]
### Define a function for comparing boolean statements for uniqueness

# %% [code]
def islogicunique(minterms1,minterms2,variables):
    """Tests whether the logic represented by minterms1 and minterms2 are functionally unique or 
    equivalent by checking all possible mappings of variables into minterms1.
    Returns [False,mapping] if a variable mapping exists that reproduces minterms2 from minterms1.
    Returns [True] if no mapping exists."""
    
    # if minterms1 and minterms2 have different numbers of minterms, then no mapping
    # between the two will exist
    if not (len(minterms1) == len(minterms2)):
        return [True]
    
    # generate all permutations (orderings) of variables
    var_permut = itertools.permutations(variables,len(variables))
    
    # generate list of dicts containing each possible permutation mapping
    all_mappings = [{variables[0]:var1,variables[1]:var2,variables[2]:var3} 
                    for (var1,var2,var3) in var_permut]
    
    # iterate through each possible mapping, checking whether it 
    # reproduces minterms1 from minterms2
    for mapping in all_mappings:
        
        # if mapping exists between minterms1 and minterms2, then they are 
        # not functionally unique
        if checkmapping(minterms1,minterms2,variables,mapping):
            return [False,mapping]
        
    return [True]

# %% [code]
def checkmapping(minterms1,minterms2,variables,mapping):
    """Checks whether a given mapping of variables into minterms1
    reproduces minterms2, returns True if mapping works"""
    
    # generate ordered list of variable mapping for re-mapping minterms1 
    var_mapping = [mapping[variables[0]],mapping[variables[1]],mapping[variables[2]]]

    # produce boolean logic statement representing minterms1 re-mapped according to mapping
    minterms1_mapped_boolean_statement = SOPform(var_mapping,minterms1)

    # compare minterms1_mapped_boolean_statement to minterms2 to see if they represent
    # equivalent logic
    minterm_eq_result = is_minterm_eq(minterms2,minterms1_mapped_boolean_statement)
    
    if minterm_eq_result:
        
        # if minterms1_mapped_boolean_statement and minterms2 are equivalent, then mapping
        # accurately describes how to reproduce the logic of minterms2 from minterms1
        return True
    
    return False

# %% [code]
def is_minterm_eq(minterms,boolean_statement):
    """Compares a list of minterms to a boolean statement,
    returns False if they are different, True if they are equivalent"""
    
    # iterate through each minterm
    for minterm in minterms:
        
        # substitute values from minterm into variables in boolean_statement
        sub_dict = {L_1:minterm[0],L_2:minterm[1],L_3:minterm[2]}
        
        
        if not boolean_statement.subs(sub_dict):
            
            # if statement does not return True, then boolean_statement does not
            # contain this minterm and minterms is not equivalent to boolean_statement
            return False

    return True
        

# %% [markdown]
## Use islogicunique() to do pairwise comparison of all logic functions with same number of minterms

# %% [code]
# create empty list for storing results of logic function comparison
mapping = []

# iterate through all logic functions in df
for df_index1,logic1 in tqdn(df.iterrows()):
    
    # generate a reduced dataframe containing all logic functions with the 
    # same number of minterms as logic1, but not including logic1 itself
    logic1_num_minterms = logic1.num_minterms
    df_logic2 = df.query('num_minterms == @logic1_num_minterms').drop(df_index1,'index')
    
    # if there are no logic functions besides logic1 with logic1_num_minterms, then
    # logic1 is unique
    if df_logic2.empty:
        mapping.append('UNIQUE')
        continue
    
    # iterate through df_logic2, comparing each logic2 function to logic1
    for sub_index,(df_index2,logic2) in enumerate(df_logic2.iterrows()):
        
        # test whether logic1 and logic2 are unique
        unique_result = islogicunique(logic1.minterms,logic2.minterms,variable_list)
        
        # if logic1 and logic2 are not unique and logic1 is not the first logic encountered
        # of a functionally equivalent set, store which logic function logic1 maps to and 
        # the mapping. This is to ensure 1 logic function of a functionally equivalent set of 
        # logic functions is stored.
        if df_index1>df_index2 and not unique_result[0]:
            mapping.append((df_index2,unique_result[1]))
            break
            
        # if logic1 has been compared to all logic2's, then logic 1 is unique.
        elif sub_index == len(df_logic2)-1:
            mapping.append('UNIQUE')


# %% [code]
# add a column to our dataframe reporting whether each logic function is unique, 
# and how it maps to another logic function if it is not unique
df = df.assign(mapping=mapping)

# %% [code]
# select combinations of minterms that are labeled unique
df_uniques = df.loc[df['mapping']=='UNIQUE']
print('The total number of functionally unique gates determined by code = '+ str(len(df_uniques)))
df_uniques

# %% [code]
# select unique combinations of minterms that contain the [0,0,0] minterm
df_uniques_zeros_on = df_uniques.query('zeros_minterm_present')
print('The total number of functionally unique gates that include the [0,0,0] minterm determined by code = '+ str(len(df_uniques_zeros_on)))
df_uniques_zeros_on

# %% [code]
df_uniques.to_csv('Functionally_unique_3input_logic_all.csv')
df_uniques_zeros_on.to_csv('Functionally_unique_3input_logic_with_zeros_minterm.csv')
```
