## Supplementary material for "Combinatorial Control through Allostery": Jupyter Notebooks where the set of functionally unique gates and constraints conditions are derived: mwc_constraints.ipynb

```
# %% [markdown]
In this notebook, we derive the necessary and sufficient set of conditions that enable us to eliminate all 3-igand responses that are incompatible with the MWC framework. As described in Appendix D.1 of the Supplementary Information, the total relative weight of the inactive states at a particular limit of ligand concentration ($\text{w}_{\text{s}^*}$) can have a multiplicative dependence on the weights at up to four other limits ($\text{w}_{\text{s}_\text{n}}$), which is expressed in a form
\begin{align}
\text{w}_{\text{s}^*} = \prod_{\text{n}=1}^4 \text{w}_{\text{s}_\text{n}}^{\alpha_\text{n}},
\end{align}
where $\alpha_n$ are real coefficients. In what follows, we first identify all possible relations of the above form and then identify the minimal set that is necessary and sufficient to account for all 3-ligand logic behaviors incompatible with MWC.

# %% [code]
# Import the required modules
import numpy as np
import itertools
np.set_printoptions(threshold=np.inf)

# %% [markdown]
First, let us define a function that outputs a string expression for $\text{w}$ at the specified limit of ligand concentrations.

# %% [code]
def toSym(arr):
    """
    Given a binary array of ligand states,
    creates a string for the corresponding weight.
    
    Examples:
    ---------
    Input: arr = [0,1,1]
    Output: 'wjk'
    
    Input: arr = [0,0,0]
    Output: 'w0'
    
    Input: arr = [1,1,1]
    Output: 'w123'
    """
    if sum(arr) == 0:
        return 'w0'
    elif sum(arr) == 3:
        return 'w123'
    
    gammaStr = ['i','j','k']
    str = 'w'
    for i in range(3):
        if arr[i]:
            str = str + gammaStr[i]
    return str

# %% [markdown]
For instance, when `arr=[0,1,1]`, corresponding to the limit where the $\text{i}^{\text{th}}$ ligand is absent while the $\text{j}^{\text{th}}$ and $\text{k}^{\text{th}}$ ligands are present at saturating concentrations, the function returns

# %% [code]
toSym([0,1,1])

# %% [markdown]
To start looking for constraint conditions, we note that each each limit $\text{w}_{\text{s}}$ has the following form:
\begin{align}
\text{w}_{\text{s}} = \omega \, \gamma_1^{\delta_1 (\text{s})} \, \gamma_2^{\delta_2 (\text{s})} \, \gamma_3^{\delta_3 (\text{s})},
\end{align}
where ${\delta_\text{i} (\text{s})}$ are binary (0 or 1) coefficients that take the value of 1 if the $\text{i}^{\text{th}}$ ligand is at a saturating concentration at the limit "s". Substituting this form into the multiplicative constraint condition, we obtain

# %% [markdown]
\begin{align}
\omega \, \gamma_1^{\delta_1 (\text{s}^*)} \, \gamma_2^{\delta_2 (\text{s}^*)} \, \gamma_3^{\delta_3 (\text{s}^*)} = \prod_{\text{n}=1}^4 \left( \omega \, \gamma_1^{\delta_1 (\text{s}_\text{n})} \, \gamma_2^{\delta_2 (\text{s}_\text{n})} \, \gamma_3^{\delta_3 (\text{s}_\text{n})} \right)^{\alpha_{\text{n}}},
\end{align}

# %% [markdown]
where $\omega = \text{e}^{-\beta \Delta \epsilon_{\text{AI}}}$. Since the algebraic condition must hold true for arbitrary choices of the MWC parameters, we equate the powers of the four MWC parameters ($\omega, \gamma_1, \gamma_2, \gamma_3$), obtaining four equations for the four unknown coefficients $\alpha_\text{n}$, namely,
\begin{align}
1 &= \sum_{\text{n}=1}^4 \alpha_\text{n}, \\
\delta_\text{k} (\text{s}^*) &= \sum_{\text{n}=1}^4 \alpha_\text{n} \delta_\text{k} (\text{s}_\text{n}), \quad 1 \le \text{k} \le 3.
\end{align}

# %% [markdown]
Choosing $\text{s}^*$ to be any of the $2^3=8$ possible ligand concentration limits, we look for all possibilities of choosing 4 other limits ($\text{s}_\text{n}$) from the remaining 7 that result in a set of real solutions for the coefficients $\alpha_\text{n}$, yielding valid constraint conditions. We then manually filter the functionally unique conditions, where none can be obtained from the other by swapping the ligand labels, yielding 8 conditions in total.

# %% [code]
# Largest number of independent limits of w
nIndep = 4

# 8 different combinations of ligand concentration limits
combinations = np.zeros([8, 3])
ind = 0
for i1 in range(2):
    for i2 in range(2):
        for i3 in range(2):
            combinations[ind] = [i1, i2, i3]
            ind += 1

for n in [2,3]: # Total number of saturating ligands
    strOutSet = []
    for i1 in range(2): # L_i is present/absent
        for i2 in range(2): # L_j is present/absent
            for i3 in range(2): # L_k is present/absent
                # Current limiting w_curr being expressed in terms of 4 others
                indCurr = np.where((combinations == [i1,i2,i3]).all(axis=1))[0][0]
                
                # Number of saturating concentration limits
                sumCurr = sum([i1, i2, i3])

                # All combinations of 4 limiting w values out of the remaining 8
                combinartionsLeft = np.delete(combinations,indCurr, axis=0)

                # Iterate over all possible combinations and see if w_curr
                # can be expressed as the product of the powers of the 4 limiting w's
                for subset in itertools.combinations(combinartionsLeft, nIndep):
                    mat = np.concatenate((np.ones([1,nIndep]), np.transpose(subset)))
                    mat = np.int64(mat)
                    if all(np.sum(subset, axis = 1) <= sumCurr): # To avoid repeat entries
                        try:
                            matInv = np.linalg.inv(mat)
                            coeffs = np.dot(matInv,np.array([1,i1,i2,i3]).transpose())
                            if sum([i1,i2,i3]) == n:
                                strOut = toSym([i1,i2,i3]) + ' = '
                                for k in range(nIndep):
                                    if coeffs[k] != 0:
                                        strOut = strOut + toSym(subset[k]) + '^' + str(coeffs[k]) + ' * '
                                strOut = strOut[0:len(strOut)-3]
                                strOutSet.append(strOut)
                        except:
                            continue

    # Remove the identical entries
    uniques = sorted(set(strOutSet))
    
    # Remove the cyclically identical entries
    # Keep wij = ... equations, but remove
    # the wik = ... ones
    if n == 2:
        uniques = uniques[0:int(len(uniques)/3)]
    uniques = sorted(uniques, key=len)
    
    # Keep only the functionally unique ones
    # (Indices manually identified;
    # to see the full list, uncomment the lines in the end)
    if n == 2:
        for i in [0, 1, 3]:
            print(uniques[i])
        print('--------------------------------------------')
    else:
        for i in [2, 3, 6, 9, 10]:
            print(uniques[i])
        print('--------------------------------------------')
    
#     # Show the full list
#     for i in range(len(uniques)):
#         print(uniques[i])
#     print('--------------------------------------------')

# %% [markdown]
Having identified the eight functionally unique constraint types, we proceed to single out those that are necessary and sufficient to eliminate all 3-ligand gates that are incompatible with the MWC framework. We write a set of auxiliary functions to perform label swapping operations, identifying the set of responses that a single condition can eliminate and updating the running list. With these function in hand, we iterate over all the subsets of 8 conditions, and write down only those that can eliminate all 152 MWC-incompatible responses, where the number 152 is the highest that any subset of constraints can eliminate.

# %% [code]
def giveResponseMat(nLimits):
    """
    Returns the list of 2**nLimits possible logic responses (0 or 1)
    at nLimits different limits of ligand concentration.
    In our case, nLimits=8 (2**3) and the number of possible
    three-input logic responses in 2**8 = 256.
    """
    nResponses = 2**nLimits
    responseMat = np.zeros([nResponses, nLimits])
    ind = 0
    for i in range(nLimits+1):
        for subset in itertools.combinations(np.arange(nLimits), i):
            responseMat[ind,list(subset)] = 1
            ind += 1
    return responseMat

# %% [code]
def giveConstraintList(leftArr, rightArr, nLimits):
    """
    Given a constraint of the form:
    \prod_i leftArr_i = \prod_j rightArr_j,
    where leftArr and rightArr are the sets of positional indices of w
    on each side of the constraint condition, the function
    returns the list of all inadmissible responses.
    
    For example, if the constraint is
    w0 x w2 x w4 = w7, then
    leftArr = [0,2,4],
    rightArr = [7],
    and all responses where the left elements and right elements
    have opposite limits (<<1 or >>1) are not realizable, regardless
    of the limits at positions [1,3,5,6].
    
    Locations: 0 1 2 3 4 5 6 7
    --------------------------
    Left:      0 - 2 - 4 - - -
    Right:     - - - - - - - 7
    --------------------------
    Free:      - 1 - 3 - 5 6 -
    
    Note that in the case of the 3-input response,
    the indicies [0,1,2,3,4,5,6,7] correspond to 
    ligand concentration limits [0,1,2,3,12,13,23,123].
    """
    
    # Number of terms in the constraint
    n = len(leftArr) + len(rightArr)
    
    # Number of free terms
    nFree = nLimits - len(leftArr) - len(rightArr)
    
    # Free indices
    indFree = np.delete(np.arange(nLimits), np.concatenate([leftArr, rightArr]))
    
    constraintMat = np.zeros([2 * 2**nFree, nLimits])
    ind = 0
    for i in range(nFree+1):
        for subset in itertools.combinations(indFree, i):
            constraintMat[ind,leftArr] = 0
            constraintMat[ind,rightArr] = 1
            constraintMat[ind,list(subset)] = 1
            ind += 1
            constraintMat[ind,leftArr] = 1
            constraintMat[ind,rightArr] = 0
            constraintMat[ind,list(subset)] = 1
            ind += 1
    return constraintMat

# %% [code]
def subtractConstraints (responseMat, constraintMat, y):
    """
    Given the list of all possible responses (responseMat),
    a vector that indicates which responses are not eliminated yet (y),
    and a list of new responses to be eliminated (constraintMat),
    the function returns the updated indicator vector (y) that reflects
    the new eliminations, as well as the total number of new eliminations (nNew).
    """
    nNew = 0
    for el in constraintMat:
        ind = np.where((responseMat == el).all(axis=1))[0][0]
        if y[ind] == 1:
            y[ind] = 0
            nNew += 1
    return [y, nNew]

# %% [code]
def iterateList(side):
    """
    Given a list of positions from the set [0,1,2,3,4,5,6,7],
    which corresponds to the ligand concentration limits [0,1,2,3,12,13,23,123],
    returns all possible lists where the labels from the original list are shuffled.
    
    Example:
    --------
    Input: [1,2,3]
    Output: [[1, 2, 3], [1, 3, 2], [2, 1, 3], [2, 3, 1], [3, 1, 2], [3, 2, 1]]
    """ 
    side = np.array(side)
    permSingle = [[1, 2, 3], \
                  [1, 3, 2], \
                  [2, 1, 3], \
                  [2, 3, 1], \
                  [3, 1, 2], \
                  [3, 2, 1]]
    permDouble = [[4, 5, 6], \
                  [5, 4, 6], \
                  [4, 6, 5], \
                  [6, 4, 5], \
                  [5, 6, 4], \
                  [6, 5, 4]]
    ind1 = np.where(side == 1)[0]
    ind2 = np.where(side == 2)[0]
    ind3 = np.where(side == 3)[0]
    ind4 = np.where(side == 4)[0]
    ind5 = np.where(side == 5)[0]
    ind6 = np.where(side == 6)[0]
    
    sideList = []
    for i in range(len(permSingle)):
        sideNew = side
        sideNew[ind1] = permSingle[i][0]
        sideNew[ind2] = permSingle[i][1]
        sideNew[ind3] = permSingle[i][2]
        sideNew[ind4] = permDouble[i][0]
        sideNew[ind5] = permDouble[i][1]
        sideNew[ind6] = permDouble[i][2]
        sideList.append(list(sideNew))
        
    return sideList

# %% [code]
def iterateListConstraint(leftArr, rightArr):
    """
    Given a list of elements on both sides of the constraint ([0,7] range),
    returns the set of functionally identical lists resulting
    from shuffling the ligand labels.
    
    Note the correspondence between indices and ligand concentration limits:
    indices: [0,1,2,3,4,5,6,7]
    limits: [0,1,2,3,12,13,23,123]
    
    Examples
    --------
    Input: leftArr = [1,2]
           rightArr = [4,5]
    Output: [[[1, 2], [1, 3], [1, 2], [2, 3], [1, 3], [2, 3]],
             [[4, 5], [4, 5], [4, 6], [4, 6], [5, 6], [5, 6]]]
    """
    leftIter = iterateList(leftArr)
    rightIter = iterateList(rightArr)
    nIter = len(leftIter)
    
    leftUnique = []
    rightUnique = []
    
    leftRightUnique = []
    for i in range(nIter):
        leftSorted = sorted(leftIter[i])
        rightSorted = sorted(rightIter[i])
        leftRightSorted = np.concatenate([leftSorted, rightSorted])
        if len(leftRightUnique) == 0:
            leftRightUnique.append(list(leftRightSorted))
            leftUnique.append(leftSorted)
            rightUnique.append(rightSorted)
        elif len(np.where((np.array(leftRightUnique) == leftRightSorted).all(axis=1))[0]) == 0:
            leftRightUnique.append(list(leftRightSorted))
            leftUnique.append(leftSorted)
            rightUnique.append(rightSorted)
    return [leftUnique, rightUnique]

# %% [code]
# Number of ligand concentration limits (2**3)
nLimits = 8

# Output option (Boolean variable)
# True: show the number of eliminated responses for every single subset
# False: show only the subsets that account for all 152 eliminated responses
showAll = False

# The 8 constraint conditions and their particular realizations, which then get relabeled
# wij = w0^-1.0 * wj^1.0 * wi^1.0              -> w12 * w0 = w1 * w2
# wij = wk^-1.0 * wj^1.0 * wik^1.0             -> w12 * w3 = w13 * w2
# wij = w0^1.0 * wk^-2.0 * wjk^1.0 * wik^1.0   -> w12 * w3^2 = w0 * w13 * w23

# w123 = w0^-1.0 * wk^1.0 * wij^1.0            -> w123 * w0 = w12 * w3
# w123 = wi^-1.0 * wik^1.0 * wij^1.0           -> w123 * w1 = w12 * w13
# w123 = w0^-2.0 * wk^1.0 * wj^1.0 * wi^1.0    -> w123 * w0^2 = w1 * w2 * w3
# w123 = wk^1.0 * wj^-1.0 * wi^-1.0 * wij^2.0  -> w123 * w1 * w2 = w12^2 * w3
# w123 = w0^-0.5 * wjk^0.5 * wik^0.5 * wij^0.5 -> w123^2 * w0 = w12 * w13 * w23

# Positional indices:   [0, 1, 2, 3,  4,  5,  6,   7]
# Concentration limits: [0, 1, 2, 3, 12, 13, 23, 123]

# List of indices (concentration limits) on the left side of constraint conditions
lefts = [[0, 4], \
         [3, 4], \
         [4, 3], \
         [7, 0], \
         [7, 1], \
         [7, 0], \
         [7, 2, 1], \
         [7, 0]]

# List of limits (positional indices) on the right side of constraint conditions
rights = [[1, 2], \
          [2, 5], \
          [0, 5, 6], \
          [3, 4], \
          [4, 5], \
          [3, 2, 1], \
          [3, 4], \
          [4, 5, 6]]

# Total number of constraint conditions
nConditions = len(lefts)

# The total number of MWC constraints
nConstraintsMax = 152

subsetMax = []

for nSet in np.arange(nConditions):
    for subset in itertools.combinations(np.arange(nConditions), nSet):
        # Total number of eliminated responses
        nConstraintTotal = 0

        # Indicator vector for tracking eliminated responses
        y = np.ones(2**nLimits)

        # List of all 2**8 = 256 responses
        responseMat = giveResponseMat(nLimits)

        # Iterate over the subset of constraint conditions
        for i in list(subset):
            # List of all possible iterations of the given constraint
            [leftUnique, rightUnique] = iterateListConstraint(lefts[i], rights[i])

            for j in range(len(leftUnique)):
                constraintMat = giveConstraintList(leftUnique[j],rightUnique[j],nLimits)
                [y, nNew] = subtractConstraints(responseMat, constraintMat, y)
                nConstraintTotal += nNew

        if showAll:
            # Show the number of eliminated responses for every single constraint
            print(str(subset) + ': ' + str(nConstraintTotal))
        else:
            # Show the subset if it accounts for the 152 eliminated responses
            if nConstraintTotal == nConstraintsMax:
                print(subset)

# %% [markdown]
As we can see, the subsets that account for all 152 eliminated 3-ligand responses contain the constraint conditions (0,1,3,4), which are
\begin{align}
\text{w}_{\text{ij}} \times \text{w}_{\text{0}} &= \text{w}_{\text{i}} \times \text{w}_{\text{j}}, \\
\text{w}_{\text{ij}} \times \text{w}_{\text{k}} &= \text{w}_{\text{ik}} \times \text{w}_{\text{j}}, \\
\text{w}_{\text{123}} \times \text{w}_{\text{0}} &= \text{w}_{\text{ij}} \times \text{w}_{\text{k}}, \\
\text{w}_{\text{123}} \times \text{w}_{\text{j}} &= \text{w}_{\text{ij}} \times \text{w}_{\text{jk}},
\end{align}
demonstrating that it is a necessary and sufficient set.

# %% [code]
```
